# Unravelling the genomic and functional arsenal of *Bacilli* endophytes from plants with different lifestyles

**DOI:** 10.64898/2026.02.06.704400

**Authors:** Nikolaos P. Arapitsas, Christos A. Christakis, Savvas Paragkamian, Stefanos Soultatos, Franziska Reden, Chrysianna Psarologaki, Emmanouil Avramakis, Alexandros Stamatakis, Emmanouil A. Markakis, Panagiotis F. Sarris

## Abstract

Endophytic microbiomes of <u>c</u>rop <u>w</u>ild relatives (CWRs) adapted to extreme environments, such as halophytes, are promising sources of plant-beneficial bacteria and secondary metabolites for sustainable food production. Here, we analyzed 25 *Bacilli* isolates obtained from CWRs, halophytes, and other plant species in Crete, Greece. Using a hybrid Illumina–PacBio sequencing approach, we generated high-quality genomes and performed comparative genomics, phylogenetic, and pangenome analyses, complemented by *in vitro* assays. We identified 312 <u>b</u>iosynthetic <u>g</u>ene <u>c</u>lusters (BGCs), nearly 60% of which showed no similarity to known clusters, revealing extensive unexplored biosynthetic potential. These unique BGCs may constitute an adaptive feature enabling endophytic *Bacilli* to colonize and interact with host plants. The isolates spanned diverse genera (*Bacillus*, *Paenibacillus*, *Peribacillus*, *Neobacillus*, *Cytobacillus*, *Rossellomorea*), including three novel species. Phenotypic assays of our isolates demonstrated high salinity tolerance (up to 17.5% w/v NaCl) and strong antagonism against major bacterial and fungal phytopathogens. Genome mining further revealed a broad array of putatively plant-beneficial traits related to growth promotion, stress adaptation, host interaction and inhibition of pathogens. Together, these findings show that *Bacilli* endophytes from wild and halophytic plants possess exceptional phylogenetic novelty, functional diversity, and biosynthetic capacity, providing new genomic and ecological insights into *Bacilli* associated with plants inhabiting extreme environments.

## Introduction

Climate change and unsustainable agricultural practices pose growing threats to food security by driving drought, soil degradation and reduced crop yields [1–4]. Plant-associated microbiomes, particularly endophytes residing within plant tissues, play key roles in host adaptation, stress tolerance, and interspecies interactions, yet their diversity and functional capacities remain underexplored in extreme habitats [5–8]. <u>C</u>rop <u>w</u>ild relatives (CWRs) and halophytes, which naturally thrive under saline or arid conditions, represent important ecological systems for studying microbial adaptation and co-evolution with plant hosts [9–11].

Members of the class *Bacilli* are ubiquitous in plant microbiomes and have evolved diverse ecological strategies to survive in fluctuating environments, including spore formation, antimicrobial production, and competitive interactions with other microbes producing diverse bioactive compounds [12–16]. Taxonomic frameworks for *Bacilli* have been repeatedly revised through 16S rRNA gene sequencing and comparative genomics, revealing previously unrecognized phylogenetic lineages and novel genera [17–23]. Despite this progress, significant gaps remain regarding their functional potential, particularly the repertoire of <u>b</u>iosynthetic <u>g</u>ene <u>c</u>lusters (BGCs) and traits mediating plant colonization and stress adaptation [24, 25].

Here, we conduct an extensive genome analysis of 25 *Bacilli* endophytes, isolated from diverse CWRs and halophytes across Crete and Chrysi islands, Greece. By integrating hybrid Illumina–PacBio genome sequencing, comparative genomics, and phenotypic assays, we examine their taxonomy, biosynthetic diversity, and potential plant-beneficial traits. Our study uncovers both the genomic novelty of these microbes and their unique functional arsenal, shedding light on the potential metabolic strategies of *Bacilli* endophytes in extreme plant-associated habitats. By characterizing genomic and functional features of endophytes associated with hosts living in extreme environments, we provide endophyte-oriented genomic insights into plant–microbe interactions and establish a foundation for future studies leveraging novel, naturally adapted microbes and microbial traits that could be tested for their effectiveness in putative applications aimed at enhancing crop resilience and supporting sustainable agricultural practices.

## Materials and Methods

### Samplings

During a three-year period (2018 - 2021) nine samplings were performed across Crete and Chrysi island, for the collection of plant individuals and/or tissues (**Figure 1**). Plant habitats included olive groves, sandy beaches, and coastal shrublands; hosts comprised cultivated species (e.g., olive trees) and CWR, with emphasis on halophytes from high-salinity areas. Sample details are provided in (Table S1). Samples were collected with sterile tools, bagged separately, and transported at 4°C to the laboratory for processing.

**Figure 1.**
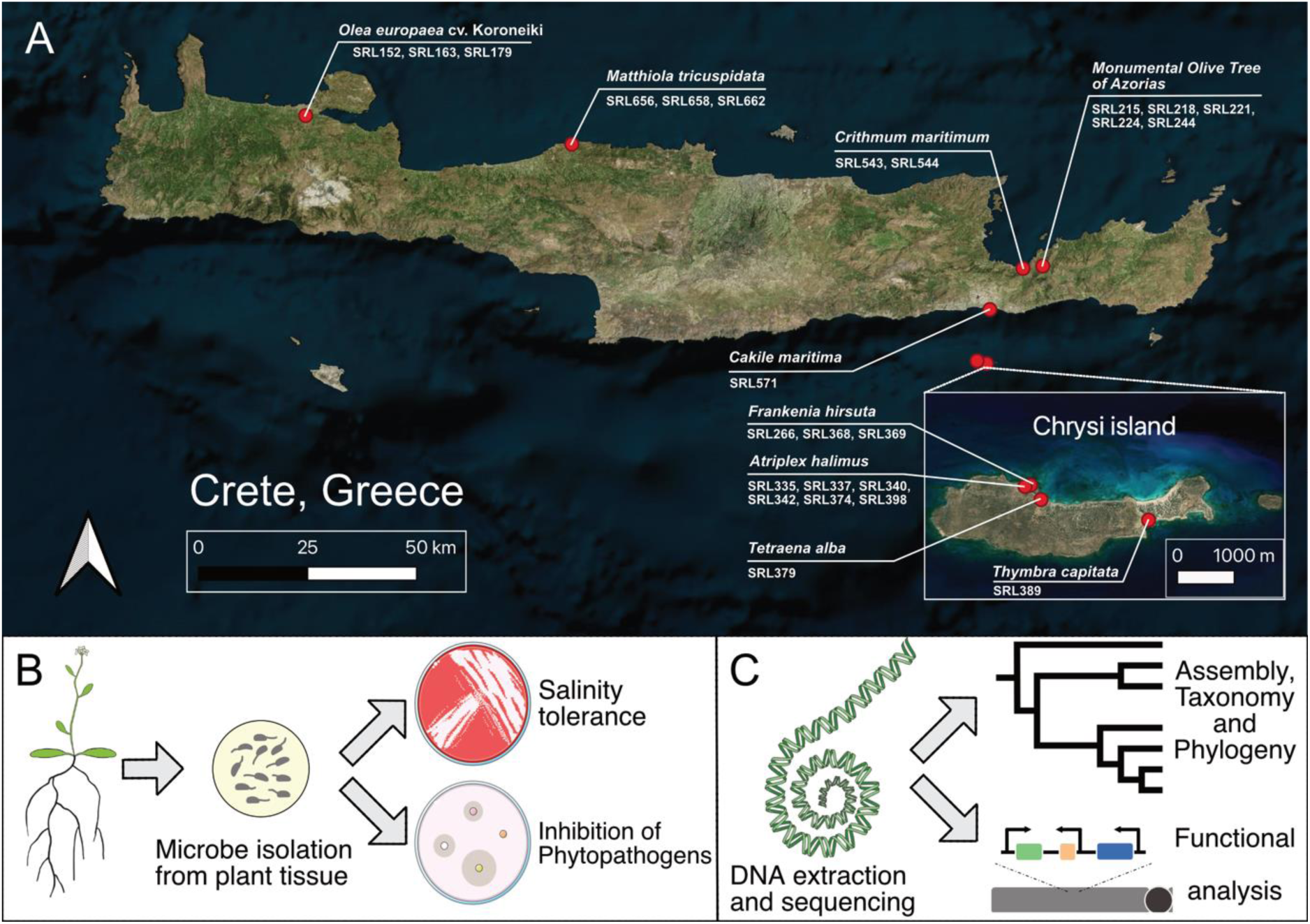
The three components of the project. **A.** Field sampling from different crop wild relative plants across Crete and Chrysi islands. The plant species and the microbial endophytes ID is displayed. **B.** The experimental procedure starts from plant tissue sterilization and microbe isolation. Then each microbe is tested against multiple phytopathogens and its maximum salinity tolerance is evaluated. **C.** DNA is extracted and sequenced using hybrid technology to investigate the genome, the taxonomy and phylogeny of each isolate. Functional and pangenome analysis is subsequently conveyed. Icons are downloaded from Bioicons except from the dendrogram which is created by Leslie Coonrod from the Noun Project.

### Microbial isolation, identification and characterization

#### Plant surface sterilization and endophytic bacterial isolation and identification

Endophytic *Bacilli* were isolated from surface-sterilized roots and leaves of selected plants based on a previously described method [9], plated on three different media (NA, R2A, NA1/2), and incubated at 28°C. Pure cultures were preserved in 50% glycerol at −80°C and preliminary identified by 16S rRNA gene sequencing. A detailed description of the sterilization, isolation, plating and culturing protocol, as well as the 16S rRNA gene-based identification of the isolates is provided in (SM 1).

#### Salt tolerance assay

The salt tolerance of each isolate was assessed by monitoring growth on NA with increasing NaCl concentrations (0.5%, 2.5%, 5%, 7.5%, 10%, 12.5%, 15%, and 17.5% w/v). Inocula from overnight cultures were streaked onto the plates and incubated at 28◦C. Growth was evaluated at 24 – 96 h post-inoculation.

#### Growth inhibition of phytopathogens

The antibacterial activity of the isolates was evaluated against four phytopathogenic bacteria: *Ralstonia solanacearum* GMI1000, *Clavibacter michiganensis* HMU4521, *Paracidovorax citrulli* ATCC29625 and *Xanthomonas campestris* pv. *campestris* strain 8004 [26]. Pathogens were grown in Nutrient Broth (NA without agar) at 28°C, 200 rpm for 48 h. Pathogen lawns were prepared by spreading cultures onto NA plates using sterile cotton swabs and *Bacilli* isolates were spot-inoculated on the same plates and incubated at 28 °C. Zones of inhibition (clear areas around the colonies of the isolates) were measured in millimeters (mm) at 72 - 96 h after incubation. Three biological replicates were performed for each isolate against each bacterial phytopathogen.

The antifungal activity of the isolates was assessed *in vitro* against four phytopathogenic fungi: *Alternaria* sp., *Botrytis cinerea*, *Fusarium oxysporum* f.sp. *radicis*-*cucumerinum,* and *Verticillium dahliae*. The isolates were tested in five biological replicates per fungus in dual-culture (confrontation test) and dual-plate (volatile test) assays following the methodology described in [9]. The effect of bacterial isolates on mycelial radial growth (mm), conidia production and hyphae width (μm), and/or (micro)sclerotia formation was evaluated and measured. For the radial growth, growth rates of fungal cultures were calculated as the mean of sequential differences in colony diameter over time per plate. Specifically, the growth step was calculated as the difference in culture diameter over the corresponding day post inoculation (DPI) interval, and the mean rate per plate was derived by averaging over all steps. Microsclerotial area (cm^2^) of *V. dahliae* was estimated by scanning the underside of each plate at the end of the bioassays and determining the black pigment area using the ImageJ 1.64r software [27]. The sclerotia number of *B. cinerea* per plate was also measured. All fungal parameters were calculated in relation to the respective controls (each fungus alone). The percentage change was calculated using the formula: Change % = [(Value_t_ - Value_c_)/Value_c_] × 100%, where Value_t_ is the treatment mean and Value_c_ is the corresponding control mean of the batch. To assess the significance of microbial effects, one-way analysis of variance (ANOVA) was conducted. *Post hoc* comparisons were performed using *Tukey’s HSD* test. Significant microbial effects, defined by a negative percent change and significant ANOVA and *Tukey* p*-*values (*p* < 0.05). Calculations were performed in R (version 4.4) with the ggplot2 package [28–30]. The raw data of the *in vitro* bioassays to evaluate the antifungal activity of the isolates are available in (SM 2).

### DNA extraction for Whole Genome Sequencing

Genomic DNA was extracted following a procedure (see: SM 1) based on a publicly available protocol by A. Kiledal and J. A. Maresca [31], modified from a method by T. Wecke. DNA samples were sequenced by Novogene (Netherlands) using both Illumina NovaSeq (2 × 150 bp) and PacBio Sequel II platforms. Information about the sequencing data is available in (SM 3). The sequences of the 25 isolates were uploaded to the European Nucleotide Archive under the PRJEB93985 study accession number. The metadata of the samples follow the MIxS-host-associated checklist of the Genomic Standards Consortium (SM 4) [32].

### Genome assembly and annotation

Illumina short reads were quality controlled with fastp v0.23.4 [33], using default settings. PacBio long reads were length checked with Filtlong v0.2.1 (https://github.com/rrwick/Filtlong) using settings “--min_length 1000” and “--keep_percent 95”. Hybrid assemblies were performed with Unicycler v0.5.0 [34] using default settings. For SRL389, SRL543 and SRL662, a first assembly was produced using flye v.2.9.2-b1786 [35]. Then, this assembly was used in Unicycler with the --existing_long_read_assembly option, using both the Illumina and PacBio processed reads. The structural statistics and functional completeness of the assemblies were evaluated using QUAST v5.0.2 [36] with default options and BUSCO 5.8.0 [37] with the *Firmicutes* database (firmicutes_odb10), respectively (see SM 5). Part of the assemblies were performed on IMBBC-HCMR’s High Performance Computing Cluster Zorba [38].

Genome annotations and genome maps (Fig. S1) were generated using Bakta v1.11.2 with its full database v6.0 [39]. Plasmid detection was performed with RFPlasmid v0.0.18 using the parameter --species Bacillus [40]. Contigs were considered as plasmids if they were predicted with high confidence by RFPlasmid and confirmed to be circular (SM 5). The predicted proteomes of the isolates were obtained from the Bakta output and used as input for OrthoFinder v.2.5.5 with default options [41].

### Taxonomic and phylogenetic analyses

A species tree was inferred from the same 120 marker genes used in the GTDB-Tk pipeline [42, 43]. Unlike GTDB-Tk, which relies on a subset of sites, full-length marker genes were employed. Genome Taxonomy Database (GTDB) reference genomes, restricted to the same putative genera as the query assemblies according to GTDB taxonomy, were selected. From this set, to avoid redundancy, highly similar genomes (≥95% average sequence identity across all marker genes; see identity matrix: SM 6) were clustered, and one genome from each cluster was retained for further analysis, resulting in overall 280 genomes, including our 25 genomes and the genomes obtained from the GTDB. Next, the corresponding marker genes were retrieved from the annotated query assemblies. Protein sequences of the 120 marker genes from both query assemblies and GTDB genomes were aligned using MUSCLE v5.1 [44] and used to project gaps onto coding DNA sequences producing codon-aware alignments. Gene trees were inferred from these DNA alignments using IQ-TREE 2 v2.4.0 [45] and combined into a species tree with ASTRAL-IV v1.22.4.6 [46]. Ballistic distances—defined as the sum of branch lengths along the unique path between two tips—were calculated between each assembly and its closest putative relative, as estimated by FastANI [47]. Species delimitation was performed with mPTP v0.2.5 [48] under maximum likelihood optimization in single-rate mode. Phylogenetic trees were visualized using TreeViewer [49]. All analyses were executed through a Snakemake workflow available on GitHub (https://github.com/FranziskaReden/bacillus_project_phylogeny.git) under the Creative Commons license (CC0 1.0) and described in more detail in (SM 1).

### Comparative Genomics and Functional Profiling

#### Pangenome analysis of selected isolates

The three putative new species and SRL368 (increased genetic novelty), were selected for pangenome analysis along with their closest relatives identified via the identity matrix from taxonomic and phylogenetic analyses (SM 6; see: Taxonomic and phylogenetic analyses). For SRL179, SRL337, and SRL543, the three closest publicly available relatives were selected; for SRL543, SRL369 — identified as a close relative — was also included. For SRL368, five public genomes were selected along with our *B. thuringiensis* isolates SRL215, SRL218, and SRL224. Genomes of the publicly available closest relatives were retrieved from NCBI using the command ncbi-genome-download bacteria with the flags --formats fasta and --flat-output, providing the desired accessions via the --assembly-accessions option [50, 51]. The pangenome analysis was made using anvi’o v8 [52, 53]. Genomic and functional annotation was conducted on genomes storage databases generated with anvi-gen-genomes-storage using anvi-run-hmms (with the flags: -I Bacteria_71 and --also-scan-trnas) [54], anvi-run-ncbi-cogs [55], anvi-run-kegg-kofams [56], anvi-run-pfams [57] and anvi-run-scg-taxonomy [58, 59]. The anvi-pan-genome command was used with --minbit 0.5, --mcl-inflation 10, and --use-ncbi-blast [52, 53]. Gene cluster presence/absence data were extracted from the SQLite PAN.db files of each pangenome analysis using sqlite3 (version 3.45.3), and post-processed with gawk (GNU Awk 5.3.0) to count gene cluster occurrences per genome. The graphs were created in R (version 4.5.1) [28] using tidyverse v2.0.0 [30], and UpSetR v1.4.0 [60].

#### BGCs, Manual curation and Genome Mining

All assemblies were uploaded to the antiSMASH version 8.0.1 platform for BGC prediction [61]. The detection strictness was set “relaxed”, and all additional features were enabled. Information about the BGC groups of each isolate and their similarity confidence with known BGCs was manually collected and groups were organized into categories (see SM 1).

Annotations were also retrieved from IMG/MER [62] and manually queried for gene terms, names, functions, products, and mechanisms for <u>P</u>lant <u>G</u>rowth <u>P</u>romotion (PGP) characteristics. To ensure annotation fidelity, annotations were kept only when annotation information was identical in at least 3 of the following database outputs: KEGG orthology [63], pfam [57], COG [55], SMART [64], Tigrfam [65], SUPERFamily [66], CATH FunFam [67]. Genome annotations were then assigned to the following categories based on KEGG orthologies and literature: “Salt Stress Tolerance”, “Disease Resistance”, “Antioxidants”, “Colonization”, “Plant Hormone & VOCs Production”, “PGP & Nutrient Acquisition”, “Chemotaxis & Flagella”, “Osmoprotection”, and “Secretion” [68–73].

PGP-related genes and gene products were derived from experimentally supported studies. Specifically, info for genes related to PGP under salt tolerance, antioxidants, and resistance to oxidative stress were derived from: [69, 70]; genes for Disease Tolerance, Plant Hormones and <u>V</u>olatile <u>O</u>rganic <u>C</u>ompound (VOCs): [68, 70–72, 74]; Osmoprotection: [69, 70]; general PGP and increased Nutrient Acquisition: [70–72]; Plant Colonization and Chemotaxis: [68, 69]; and Secretion: [69, 73]. An annotation was accepted only if two or more systems supported the same function. Detailed gene lists and references are provided in SM 7.

Gene counts across functional categories and isolates were visualized using R (v4.5.1) [28] with the pheatmap package (v1.0.13) [75], while BGC plots were created using ggplot2 v3.4.4 [29], tidyverse v2.0.0 [30], patchwork v1.3.1 [76], and cowplot v1.1.3 [77].

## Results and Discussion

### Genomic Characterization and Taxonomic Diversity

A total of 25 endophytic *Bacilli* were isolated from eight plant species across nine sampling sites in Crete and the nearby Chrysi satellite island (**Figure 1**; **Table 1**; Table S1). Twenty isolates were originated from root samples and 5 from leaves (**Table 1**). Thus, reflecting a broad environmental sampling from olive trees, halotolerant plants and halophytes from different locations, tissues and diverse lifestyles.

**Table 1.**
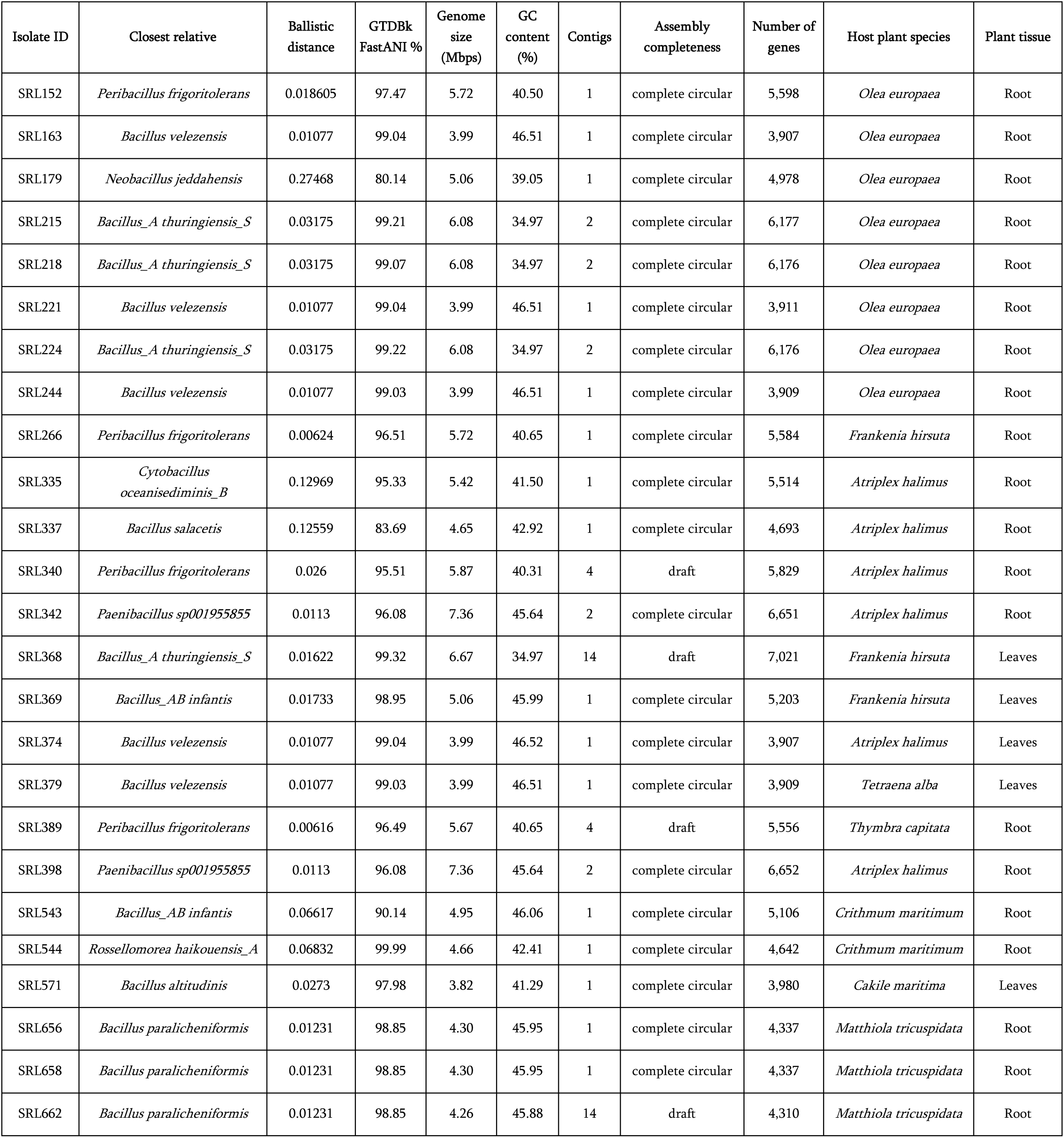
Selected features of the 25 *Bacilli* isolates and their assemblies. Gene numbers are obtained from Bakta annotation. For SRL368, SRL389 and SRL662, the contig number refers to contigs bigger than 1000 bp. A more comprehensive version of this table, including additional info for every isolate, is available in **SM 5**.

The hybrid sequencing strategy enabled high-quality assemblies with 21 out of 25 being assembled into complete circular genomes. The isolates SRL340, SRL368, SRL389, and SRL662, were draft assemblies, consisting of 2–14 contigs (≥1 kb) (**Table 1**; SM 5). Genome sizes ranged from 3.82 Mb (SRL571; *Bacillus altitudinis*) to 7.36 Mb (SRL342 and SRL398 assigned as *Paenibacillus sp001955855*), with a mean genome size of approximately 5.16 Mb (**Table 1**; SM 5). The genus *Bacillus* was the most prevalent (16 isolates), followed by *Peribacillus* (4), *Paenibacillus* (2), and one isolate each from the genera *Neobacillus*, *Cytobacillus*, and *Rossellomorea* (**Table 1**; SM 5).

The GC content spanned from 34.97% (SRL215, SRL218, SRL224, and SRL368; all *Bacillus thuringiensis*) to 46.52% (SRL374, *Bacillus velezensis*), averaging 42.51% across the dataset (**Table 1**; SM 5). The number of predicted genes, based on Bakta annotation, ranged from 3,907 (SRL163, *B. velezensis*) to 7,021 (SRL368, *Bacillus thuringiensis*) (**Table 1**; SM 5). The diversity in genome size and features within closely related *Bacilli* species underscores potential genomic adaptations to host plant species or environmental niches, as well as traits possibly acquired through horizontal gene transfer [78, 79]. For example, *Bacillus thuringiensis* SRL368 exhibited a notably large genome (6.67 Mb) and high gene content (7,021), which may be influenced by plasmid content and isolate-specific genomic expansions (**Table 1**; SM 5). Four putative plasmids were identified in SRL368; however, its fragmented assembly may have led to an underestimation (SM 5).

### Phylogenetic analysis of *Bacilli* class including the 25 isolates

To explore the taxonomic diversity and placement among the assembled genomes, a species tree was constructed based on 120 GTDB marker genes (**Figure 2**) [42, 43]. Although not intended as a definitive reconstruction of evolutionary history, the tree was used to visualize genome relationships and complement genome-wide similarity estimates such as those derived from FastANI (**Table 1**) [47]. The topology of the tree generally reflects the established GTDB taxonomy (**Figure 2**), with most assemblies clustering according to their genus-level classifications, some of which are, notably, GTDB-specific [43]. Ballistic distances calculated on the tree are largely consistent with FastANI values, suggesting that divergence in marker genes captures broader genomic similarity patterns reasonably well (**Table 1**). Taxonomic authority of the genus *Bacillus* is ambiguous and GTDB has the most comprehensive phylogeny. For example, *Bacillus_AB* is closer to *Cytobacillus* than to *Bacillus* and it does not exist in the NCBI taxonomy.

**Figure 2.**
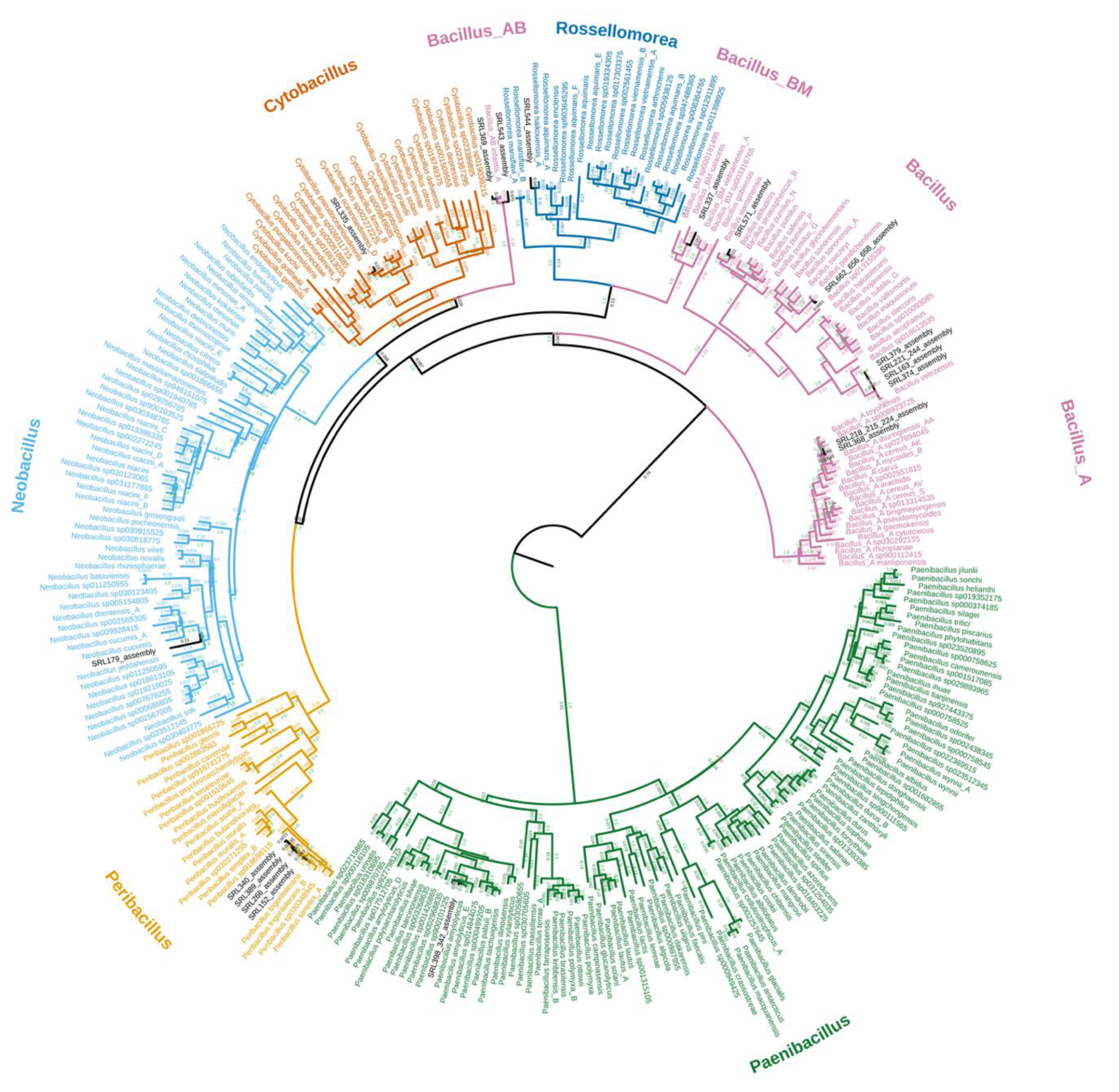
Phylogenetic tree including the 25 endophytic *Bacilli.* Phylogenetic tree showing the placement of the 25 endophytic *Bacilli* assemblies (black font) among the publicly available reference genomes. Numbers above the internal branches (color-coded from green to red according to their value) indicate branch support values (local posterior probabilities). Numbers below the branches represent branch lengths in coalescent units. The headers display the genera of the strains which share the same color.

Three isolates, SRL179 (*Neobacillus* sp.), SRL337 (*Bacillus* sp.), and SRL543 (*Bacillus* sp.), exhibited low FastANI values (80.14%, 83.69%, and 90.14%, respectively) compared to their closest reference genomes (**Table 1**). They also appear to have substantially diverged from their closest relatives in the inferred phylogenetic tree, with ballistic distances of 0.27468, 0.12559, and 0.06617, respectively, providing strong evidence that they may represent novel species (**Table 1**; **Figure 2**). Consistent with these findings, species delimitation with mPTP [48] estimated SRL179, SRL337, and SRL544 to be unique species, providing additional evidence that at least two of these lineages likely represent previously undescribed taxa.

While the species tree was primarily used to assess taxonomic placement and for visualization, it provides useful insights into the diversity and divergence present within the dataset. To better resolve evolutionary relationships within *Bacillota* genera, future research could explore whole-genome-based inference or incorporate additional genus-specific marker sets for phylogenetic analysis.

### *In vitro* assays for salinity tolerance and growth inhibition of selected phytopathogens

#### Salinity tolerance

Plant-associated salt-tolerant bacteria can contribute to salt stress alleviation and growth promotion of crops [9, 10, 80]. Most isolates (18) grew at salinity of 7.5% w/v or higher, while all remaining isolates tolerated salinity levels of up to 5% w/v (SM 5).

The putative novel species SRL179 and SRL337 grew at salinity levels up to 17.5% w/v, whereas SRL543 tolerated up to 10% w/v salinity (SM 5). Notably, the *Peribacillus frigoritolerans* isolate SRL266 tolerated salinity as high as 17.5% w/v, while the other *Peribacillus* isolates SRL152, SRL340 tolerated up to 5% w/v and SRL389 tolerated up to 7.5% w/v salinity (SM 5). Similarly, while *Bacillus thuringiensis* SRL368 grew at salinity levels of up to 17.5% w/v, other isolates of the same species, SRL215, SRL218, and SRL224, could not tolerate salinity levels above 5% w/v (SM 5). These properties could confer an advantage to SRL266 and SRL368 for potential applications and underscore their divergence from their close phylogenetic relatives.

#### Αntibacterial activity

The 25 *Bacilli* isolates were evaluated for their ability to inhibit the economically important bacterial phytopathogens *Clavibacter michiganensis*, *Ralstonia solanacearum*, *Paracidovorax citrulli*, and *Xanthomonas campestris* pv. *campestris*. Inhibition was most prevalent against the Gram-positive *C. michiganensis*, with 11 isolates displaying clear zones of inhibition ranging from 2 to 20 mm in diameter (**Figure 3A**; SM 5). Notably, SRL656, SRL658, and SRL662 exhibited the strongest activity (**Figure 3A**; SM 5). Among the *Bacillus thuringiensis* isolates, SRL215, SRL218, and SRL224 inhibited *C. michiganensis*, whereas SRL368 showed no inhibition, again highlighting the variation in the characteristics and properties of separate *B. thuringiensis* isolates (SM 5). In contrast, the other bacterial pathogens, all being Gram-negative, were inhibited by less isolates, and the resulting zones of inhibition were minimal or barely detectable (1–2 mm) (SM 5).

**Figure 3.**
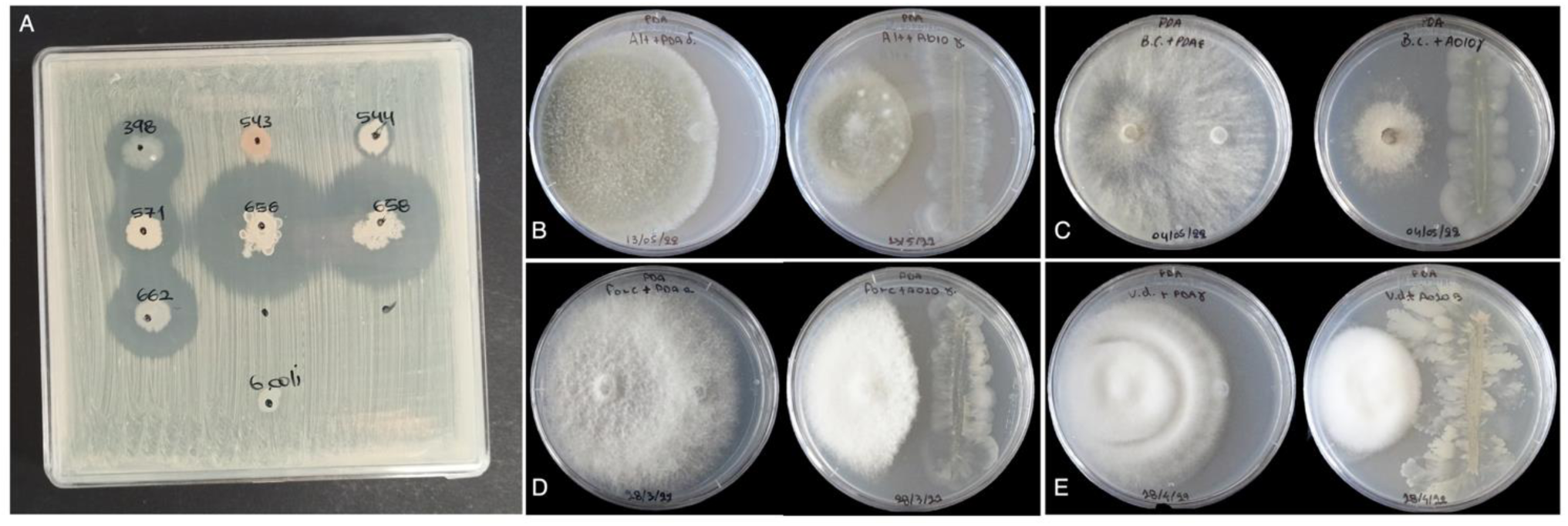
*In vitro* bioassays against selected phytopathogens. **A.** Inhibition of bacterial growth of *Clavibacter michiganensis* by the isolates SRL398, SRL571, SRL656, SRL658 and SRL662. Inhibition of fungal growth of **B.** *Alternaria* sp., **C.** *Botrytis cinerea*, **D.** *Fusarium oxysporum* f.sp. *radicis-cucumerinum* and **E.** *Verticillium dahliae* by the most effective *Bacillus* isolate (SRL163) in dual-culture assays.

#### Αntifungal activity

Based on preliminary *in vitro* screening, 15, 17, 16 and 17 bacterial strains exhibited antagonistic activity against *Alternaria* sp., *Botrytis cinerea*, *Fusarium oxysporum* f.sp. *radicis-cucumerinum* and *Verticillium dahliae,* respectively (Table S2). Thus, they were selected for further testing.

In thorough dual-culture assays, the mycelial growth of *Alternaria sp.*, *B. cinerea*, *F.o*. f.sp. *radicis-cucumerinum* and *V. dahliae* was inhibited by 12, 15, 13 and 15 isolates, while conidial production of the abovementioned fungi was reduced by 0, 12, 13 and 10 isolates, respectively (**Figure 4**; Fig. S2, Fig. S3). Moreover, the hyphae width of *B. cinerea*, *F.o*. f.sp. *radicis-cucumerinum* and *V. dahliae* was affected by 8, 12 and 5 isolates respectively, 13 isolates reduced *V. dahliae* microsclerotial area, whereas the number of *B. cinerea* sclerotia was not affected by any of the *Bacillus* isolates tested (**Figure 4**; Fig. S4). These are statistically significant findings which align with earlier studies highlighting the broad-spectrum antifungal activity of *Bacillus* spp., commonly attributed to cyclic lipopeptides (CLPs) and hydrolytic enzymes [81]. Specifically, strains of *B. pumilus* and *B. subtilis* producing surfactin, iturin and fengycin showed antimicrobial activity against several phytopathogens including *B. cinerea, F. oxysporum, R. solani, S. sclerotiorum, Pythium ultimum* and *Phytophthora capsici* [82].

**Figure 4.**
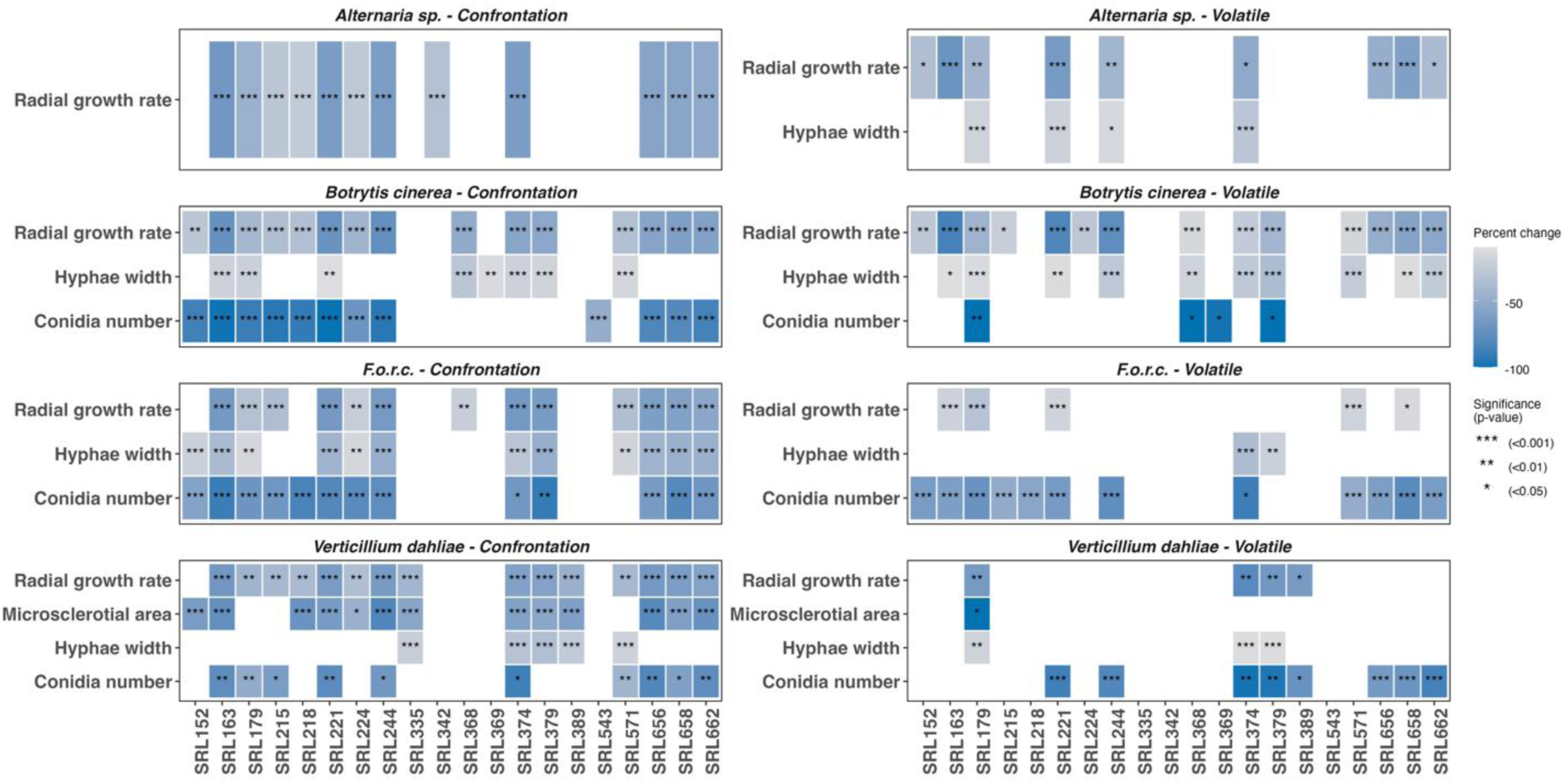
Graph of antifungal activity of bacterial isolates. Antifungal activity of bacterial isolates against four fungal plant pathogens (*Alternaria* sp., *Botrytis cinerea*, *Fusarium oxysporum* f.sp. *radicis*-*cucumerinum* and *Verticillium dahliae*) in *in vitro* assays (confrontation and volatile), estimated with four variables. The color showcases the percent change based on the control of the assay and stars indicate the statistical significance based on the Tukey HSD post-hoc test.

In thorough dual-plate assays for activity via VOCs, 9 isolates inhibited *Alternaria* sp., 14 *B. cinerea*, 5 *F. o*. f.sp. *radicis-cucumerinum* and 4 *V. dahliae* (**Figure 4**; Fig. S5). These results suggest that VOCs produced by *Bacillus* spp. also exert indirect antifungal effects against certain plant pathogenic fungal species [83]. Such VOC-inducing effects have also been reported against *B. cinerea* in previous studies [84]. VOCs that are commonly produced by *Bacillus* spp. include ketones, alcohols, aldehydes, acids, pyrazines and terpenes. Some of these compounds have been proved effective in inhibiting fungal spore germination and mycelial radial growth *in vitro* [85, 86]. Similarly, *B. velezensis* and *B. subtilis* have been shown to emit VOCs that reduce the fungal growth, sporulation and change hyphae morphology [84, 85]. In the present study, VOCs emitted by 4 *Bacillus* isolates affected conidia production of *B. cinerea*, whereas VOCs of 12 and 8 isolates affected conidial production of *F.o*. f.sp. *radicis-cucumerinum* and *V. dahliae*, respectively. However, none of the tested isolates reduced significantly conidial production of *Alternaria* sp. via VOCs-inciting effects. Hyphal width of *Alternaria* sp., *B. cinerea*, *F. o*. f.sp. *radicis-cucumerinum* and *V. dahliae* was significantly altered by 4, 10, 2 and 3 isolates, respectively (**Figure 4**; Fig. S6). Moreover, one isolate reduced significantly *V. dahliae* microsclerotial area, but none affected *B. cinerea* sclerotia number (**Figure 4**; Fig. S7).

Overall, the findings of the present study suggest significant antifungal potential of several *Bacillus* spp. isolates against major plant pathogenic fungi, through both diffusible and volatile-mediated compounds. The identification and the chemical characterization of the active compounds would be of great interest, and it remains to be addressed in future work.

### Genomics and functional analysis

#### Comparative Genomics with OrthoFinder

OrthoFinder identified 12,162 orthogroups across the 25 *Bacilli* isolates, encompassing 120,736 genes (97.1% of the 124,385 genes in total), including 804 single-copy orthogroups (Table S3). The remaining 3,649 genes (2.9%) were unassigned (not classified into any orthogroup), and 253 isolate-specific orthogroups (681 genes, 0.5%) were also detected (Table S3).

Isolate-specific orthogroups served as a measure for assessing the genetic novelty across the 25 isolates (Fig. S8A). All three putative new species - SRL179, SRL337, and SRL543 - harbored high numbers of unique orthogroups. SRL543 had 8 isolate-specific orthogroups (18 genes, 0.4% of its genes in isolate-specific orthogroups); SRL337 had 27 (104 genes, 2.3%) the third highest among all isolates; and SRL179 had 48 (113 genes, 2.3%) the second highest number of isolate-specific orthogroups (Fig. S8A; SM 8). Notably, *Bacillus thuringiensis* SRL368, exhibited the highest number of isolate-specific orthogroups: 105 orthogroups (293 genes; 4.3%) (Fig. S8A; SM 8). Despite SRL368 belonging to the same species with SRL215, SRL218, and SRL224, it harbors a markedly greater number of unique orthogroups, suggesting additional intra-species genomic divergence (Fig. S8A; SM 8).

In most cases, isolates with many isolate-specific orthogroups also showed higher proportions of unassigned genes (Fig. S8B). This trend may reflect greater genomic divergence or that certain lineage-specific genes are not captured in shared or isolate-specific orthogroups, highlighting the potential for even more undiscovered genetic novelty within *Bacilli*. In addition to 1,202 core orthogroups shared across all isolates, the number of accessory-orthogroups (shared by 2 or more isolates) varied substantially (Table S3; Fig. S8C; SM 8).

#### Pangenome analysis of the three putative novel species and SRL368 with their closest relatives

Pangenome analysis enabled comparisons of the genetic features of the three putative new species and isolate SRL368 with their closest relatives. The pangenome of SRL179 and its three closest relatives contains 10,261 gene clusters. This pangenome shows extensive divergence among SRL179 and its closest relatives, with nearly 60% of the gene clusters being species-specific, and only 21.6% constituting the core genome (**Figure 5A**). SRL179 also harbors 1,077 species-specific gene clusters absent from the other genomes, underscoring its strong genomic uniqueness (**Figure 5B**). The pronounced divergence between SRL179 and its relatives highlights the potential of this clade—and SRL179 in particular—to encompass putatively novel functional and genetic features [87].

**Figure 5.**
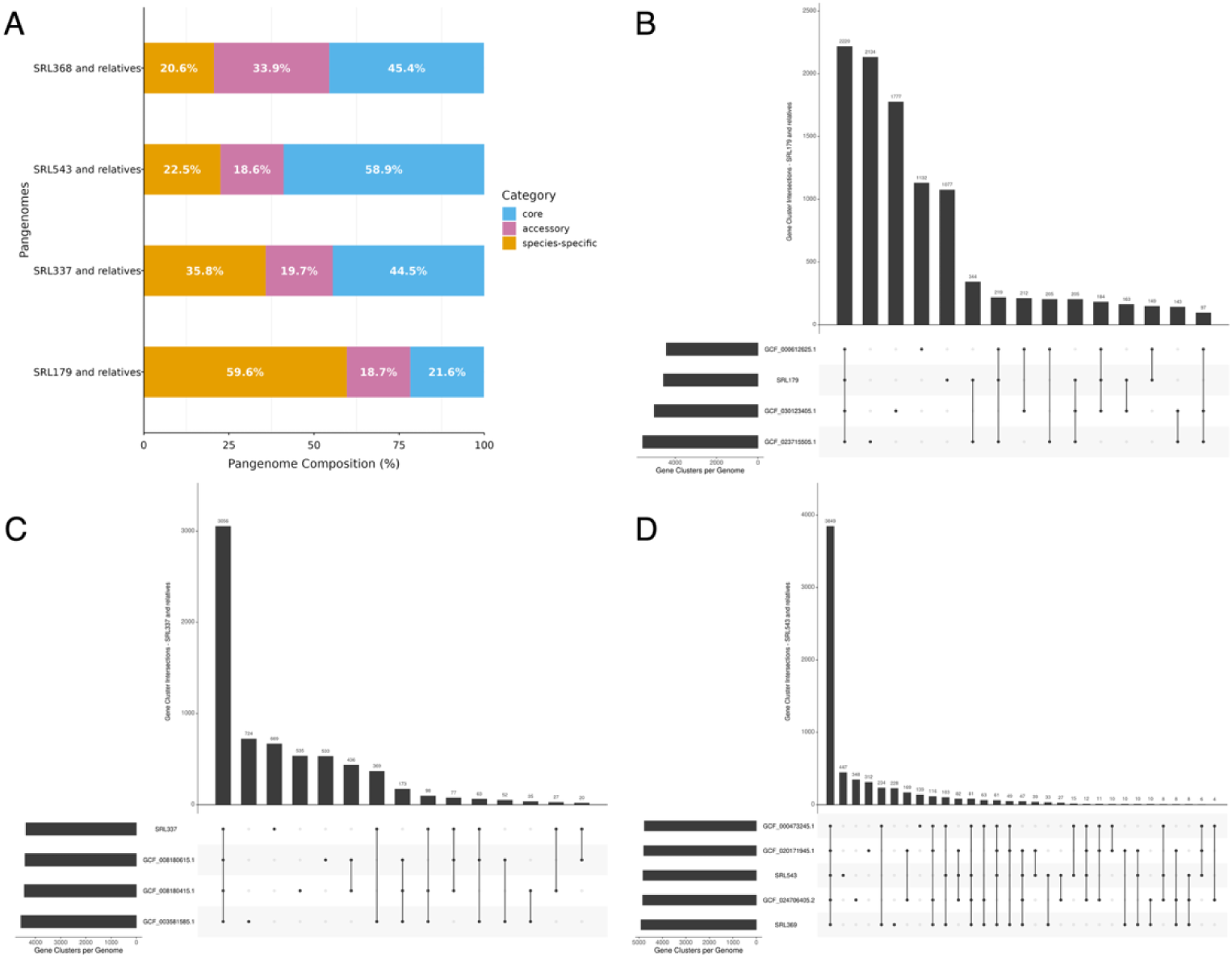
Pangenome analysis of selected isolates with their closest relatives. **A**. Distribution of core, accessory, and species-specific genes (%) in the pangenome analyses of isolates SRL179, SRL337, SRL543, and SRL368 with their closest relatives. (**B–D**) UpSet plots for the pangenome analyses of the three putative new species: **B**. SRL179, **C**. SRL337 and **D**. SRL543, with their closest relatives. In the case of SRL543, the *Bacillus infantis* isolate SRL369 was also identified as a close relative and included in the analysis. The UpSet plot for the pangenome analysis of the isolate SRL368 with its closest relatives is provided in (**Fig. S6**). Horizontal bars (left) indicate the total number of gene clusters per genome, while vertical bars (top) display the number of shared clusters for each intersection (genome combination) indicated by the connected dots below.

The pangenome of SRL337 and its three closest relatives includes 6,867 gene clusters. It also exhibits substantial divergence, with 35.8% species-specific and 44.5% core gene clusters (**Figure 5A**). SRL337 possesses 669 species-specific gene clusters, the second highest among its relatives, highlighting pronounced differences in gene content relative to its closest relatives (**Figure 5C**).

SRL543 was compared with its three closest publicly available relatives and the closely related *Bacillus infantis* SRL369. Their pangenome (6,539 gene clusters) includes 22.5% species-specific and 58.9% core genes (**Figure 5A**). Interestingly, SRL543 has the most species-specific clusters (447) among its relatives, indicating a high degree of isolate-specific gene content (**Figure 5D**).

In *Bacillus thuringiensis* SRL368, 33.9% of clusters were accessory—more than in the other pangenomes—possibly suggesting a broader distribution of certain traits shared across separate *B. thuringiensis* strains and isolates (**Figure 5A**). A total of 4,308 clusters (45.4%) formed the core genome, while 20.6% were species-specific (**Figure 5A**). Notably, SRL368 contained the highest number of gene clusters and 1,154 uniquely shared clusters (the second-largest gene cluster intersection following the core genome) with the accession GCF_020775355.1, the genome with the second-highest number of gene clusters (Fig. S9). Additionally, SRL368 contained 15 unique clusters absent from all other genomes (Fig. S9). These results indicate that SRL368 harbors novel gene clusters and features not widely distributed among most of its relatives. Meanwhile, SRL215, SRL218 and SRL224 –from the same olive tree root sample – (**Table 1**; SM 5), did not have species-specific gene clusters individually, but collectively shared 238 unique clusters absent from the other genomes (Fig. S9). These shared clusters may also encode novel features that warrant further investigation.

#### <u>B</u>iosynthetic <u>G</u>ene <u>C</u>lusters (BGC) prediction

antiSMASH identified and annotated 312 BGCs across the 25 *Bacilli* isolates (SM 9). Isolates SRL215, SLR218, SRL224 and SRL368 (*B. thuringiensis*) and SRL662 had the highest BGC number (17) (Fig. S10; SM 9). Although SRL662 has a fragmented genome (**Table 1**; SM 9), which may inflate BGC predictions, its increased BGC count compared to SRL656 and SRL658, all *B. paralicheniformis* from the same *Matthiola tricuspidata* root system, suggests that subtle genetic, functional, and biosynthetic differences can occur even within a single host microenvironment (**Table 1**; Fig. S10, SM 5, SM 9). Analogous, phenotypic and functional divergence has been reported among *B. licheniformis* isolates from the same source [88].

Mean BGC counts per genus revealed *Bacillus* and *Paenibacillus* as the most BGC-rich, followed by *Peribacillus*, and then *Neobacillus*, *Rossellomorea*, and *Cytobacillus* (Fig. S11), aligning with previous large-scale findings on *Bacillales* genomes [25, 89]. Among the eight BGC classes (SM 1), terpenes were the most frequent (23.7%) (Fig. S12; SM 9), highlighting their widespread role in the metabolite arsenal of *Bacilli* (SM 9). Known for their antimicrobial properties and potential applications against multidrug-resistant pathogens [90, 91], terpenes are naturally produced by *Bacilli* species which are promising microbial platforms for large-scale terpene production [92, 93]. The “others” class was the second most frequent (19%), underscoring broad chemical diversity in *Bacilli* isolates, while NRPS (14.7%), PKS (12.5%), and RiPPs (11.2%) were also notably represented (SM 9).

antiSMASH also assessed BGC similarity to known clusters (**Figure 6**; Fig. S10; SM 9). About 25% of the BGCs exhibited high similarity, mainly with NRPS-other hybrids, 5% medium similarity, mostly NI-siderophores (SM 9), and 11% low (mostly terpenes and betalactones) (SM 9). Notably, 59% showed no detectable similarity to any known cluster, highlighting a considerable novel biosynthetic potential (SM 9). Among these BGCs, terpenes were the most frequent, followed by NRPS and PKS clusters (SM 9), which are associated with antifungal biocontrol and plant-growth promotion [94]. These clusters underpin plant–microbe interactions and offer promising targets for further metabolic studies and biosynthetic engineering [94].

**Figure 6.**
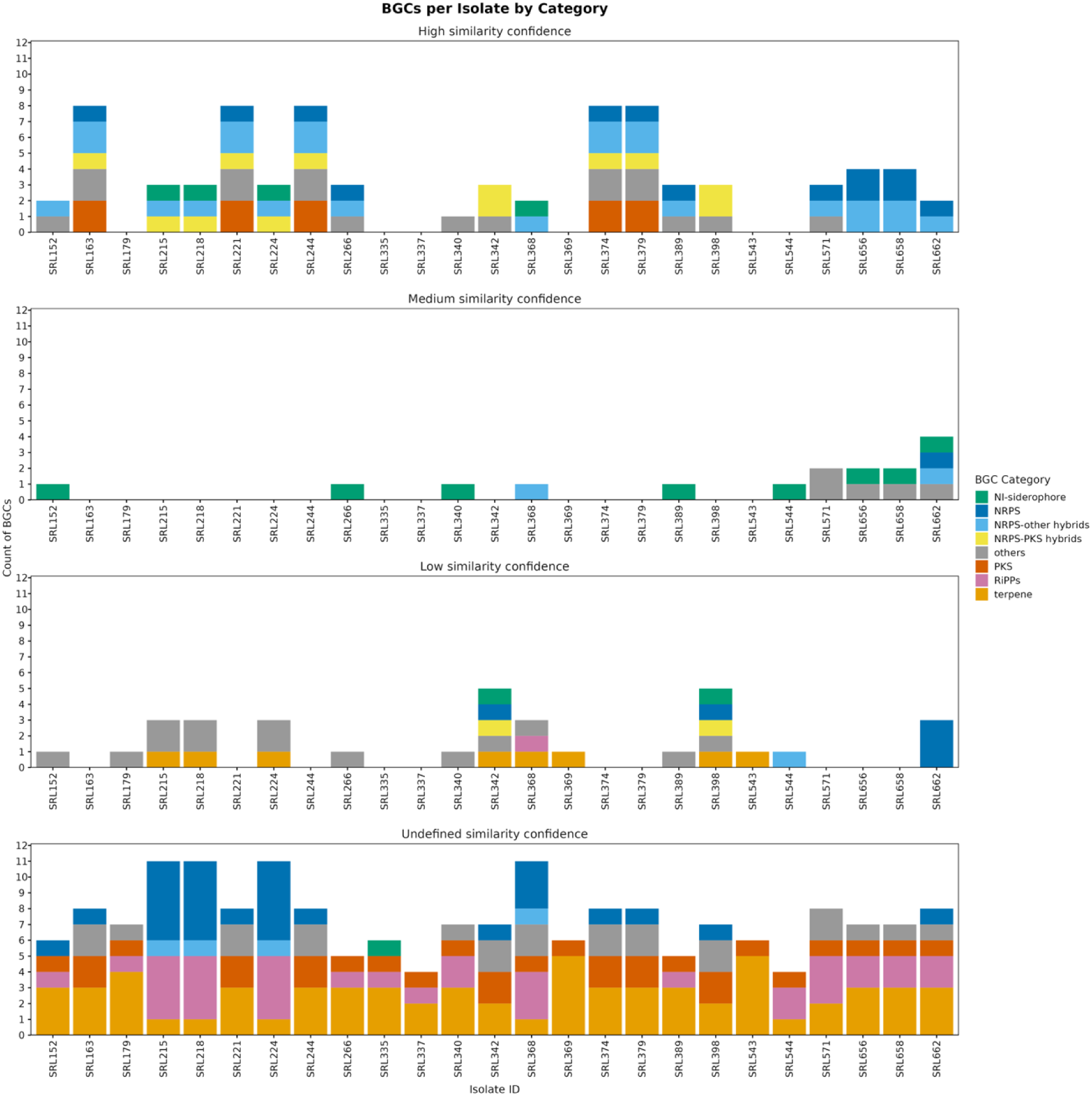
Distribution of BGCs across isolates by category and similarity confidence to known BGCs. Stacked bar plots showing the number of BGCs identified per isolate, grouped by similarity confidence levels to known BGCs (from top to bottom: High, Medium, Low, Undefined) as assigned by antiSMASH. BGCs are further grouped into eight categories: NI-siderophores, NRPS, NRPS-PKS hybrids, NRPS-other hybrids, others, PKS, RiPPs and terpenes (see SM 1 - Supplementary Methods), indicated in the plot by different colors. “Undefined similarity” refers to clusters without close matches to known BGCs in the antiSMASH database.

Comparisons between isolates of the same species revealed putative niche- or host-associated divergence and potential horizontal BGC transfer. In some cases, isolates showed highly conserved profiles. *B. velezensis* isolates (SRL163, SRL221, SRL244, SRL374, SRL379) from different hosts shared identical repertoires, enriched in terpenes, PKS, and “other” types (e.g., lanthipeptides, phosphonates). Notably, half of their clusters were of undefined similarity, highlighting novel biosynthetic potential despite any taxonomic and ecological homogeneity (**Table 1**; **Figure 6**; Fig. S10, S12; SM 9). Similarly, *B. infantis* SRL369 and SRL543, from distinct hosts, displayed identical profiles dominated by terpenes and one PKS cluster, despite SRL543 likely representing a novel species (**Table 1**; **Figure 6**; Fig. S10, S12; SM 9). All but one of their BGCs had undefined similarity, positioning *B. infantis* and especially SRL543 as a promising candidate for further study (**Figure 6**; Fig. S10; SM 9).

In contrast, *B. thuringiensis* isolates possessed both the highest total and undefined-similarity BGC counts, indicating strong biosynthetic potential (SM 9; Fig. S10). SRL215, SRL218, and SRL224 (from olive roots) shared similar repertoires, whereas SRL368 (from *Frankenia hirsuta* leaves) showed distinct BGC diversity (**Table 1**; **Figure 6**; Fig. S10; S12; SM 9). These differences may reflect host-associated structuring or isolate-specific acquisition of biosynthetic functions, a hypothesis that could be validated in the future using pipelines to detect putative horizontal gene transfer events.

The putative novel species SRL179 and SRL337 also showed exceptional BGC novelty. All BGCs in SRL337 and all but one in SRL179 lacked similarity to known clusters (**Figure 6**; Fig. S10; SM 9). Both carried one PKS- and one RiPP-associated BGC; SRL179 also harbored a phosphonate BGC of undefined similarity and a low-similarity betalactone cluster (**Figure 6**; Fig. S12; SM 9). These chemically rich and phylogenetically distinct strains are strong candidates for future biosynthetic and functional studies [24, 95].

Overall, the 25 *Bacilli* isolates revealed extensive BGC diversity and novelty, with most clusters unrelated to known BGCs. *Bacillus* and *Paenibacillus* were the most BGC-rich, with *B. thuringiensis* standing out for both abundance and the presence of numerous uncharacterized clusters, including isolate-specific variation likely shaped by ecological factors. The putative new species SRL179 and SRL337 also exhibited pronounced divergence in their biosynthetic repertoires. Together, these findings underscore the widespread presence of previously uncharacterized BGCs among endophytic *Bacilli* and highlight the potential of both well-studied and novel lineages as reservoirs of unexplored secondary metabolites.

#### Gene mining for secondary metabolites and genes involved in host-microbe interactions

The overall gene count distribution across nine functional categories revealed substantial functional diversity among the 25 isolates. Genes associated with Salt and Stress Tolerance, Chemotaxis, Disease Resistance, and Plant Colonization were the most abundant and widely distributed.

Gene mining revealed that the closely related *Bacillus thuringiensis* SRL215, SRL218, and SRL224 contained identical numbers of genes involved in host-microbe interactions and secondary metabolites, as did *B. velezensis* isolates SRL163, SRL221, SRL244 and SRL379 (**Figure 7**). Combined with the phylogeny, these data indicate that these *B. thuringiensis* isolates are likely close relative isolates of the same initial microbe originated from the same plant. While the four *B. velezensis* isolates exhibit notable similarities, they are originated from different plant samples (**Table 1**; SM 5). In contrast, *B. paralicheniformis* SRL662, from the same plant sample as SRL656 and SRL658, showed a distinct profile with markedly higher counts in secondary metabolite, chemotaxis, and salt-tolerance genes (**Figure 7**). Differences between SRL662 and SRL656/658 were also evident in their BGC profiles, as discussed above. Notably, OrthoFinder did not differentiate these isolates through isolate-specific orthogroups (Fig. S8A), emphasizing that BGC prediction and manual genome mining can uncover diversity overlooked by orthogroup analysis alone.

**Figure 7.**
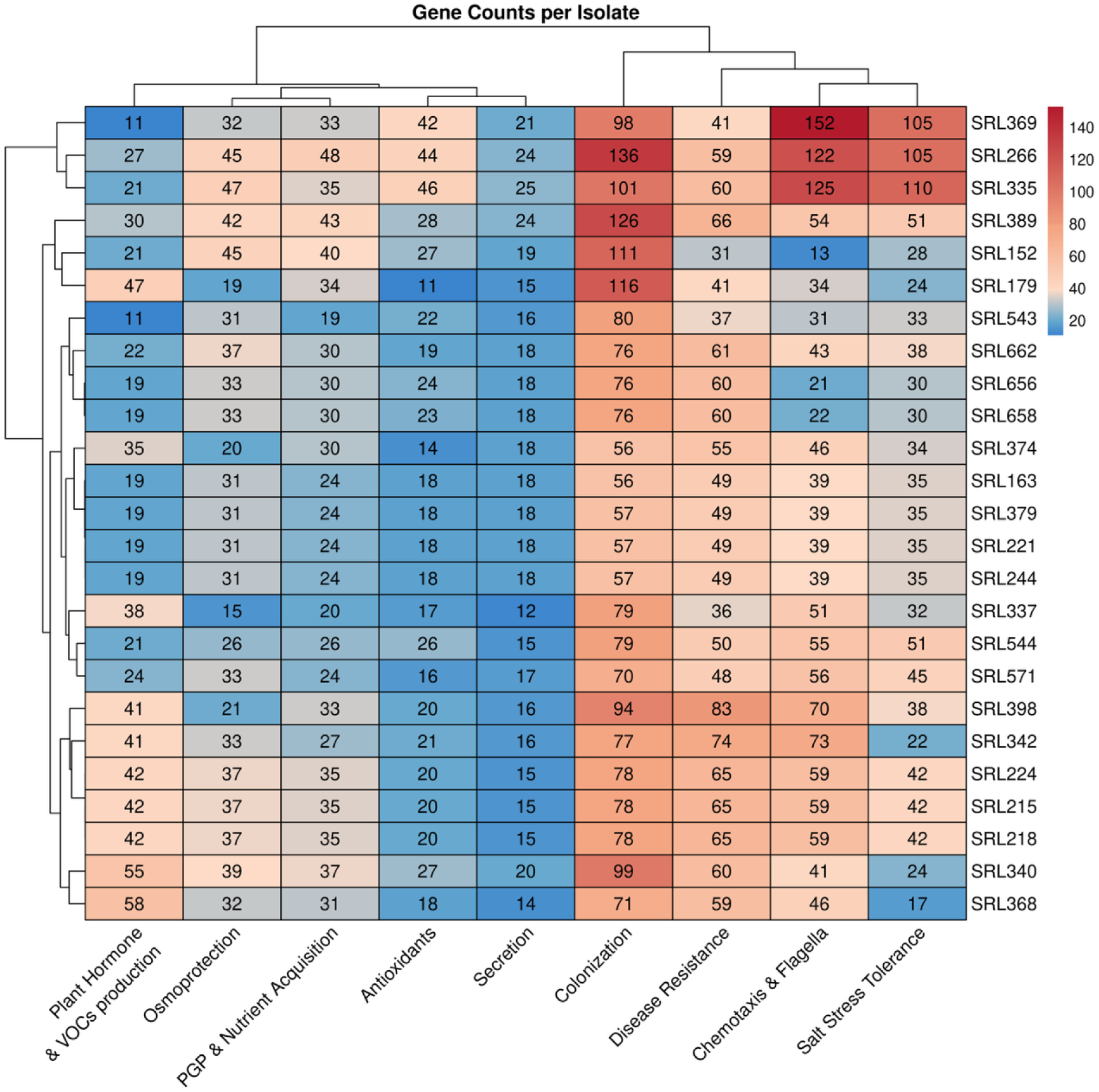
Heatmap of gene counts associated with plant–microbe interactions across the 25 bacterial isolates. Heatmap showing the distribution of gene counts across nine plant–microbe associated functional categories: Plant hormone and VOCs production, Osmoprotection, PGP and Nutrient Acquisition, Antioxidants, Secretion, Colonization, Disease Resistance, Chemotaxis and Flagella, and Salt Stress Tolerance (see: BGCs, Manual curation and Genome Mining). Rows represent isolates and columns represent functional categories. Numbers within cells indicate the gene counts, while the color gradient reflects relative abundance. Hierarchical clustering is applied to both isolates (rows) and functional categories (columns).

*Peribacillus frigoritolerans* SRL266 exhibited stronger functional potential than other isolates, including those of the same species, especially in chemotaxis, plant colonization, salt stress tolerance and PGP- and nutrient-acquisition (**Figure 7**). *B. infantis* SRL369 contained the most chemotaxis-related genes (152) and ranked among the top isolates in salt-tolerance (105) and colonization (98), suggesting strong host-sensing capacity and adaptation (**Figure 7**). Its close relative SRL543, likely a novel species, also had many colonization genes but fewer genes in other categories, highlighting a distinct functional profile. The inability of BGC profiles to differentiate these functional traits underscores the value of a more holistic approach that integrates genome mining with functional analysis. Among all isolates, *Cytobacillus* SRL335 contained the highest Osmoprotection, Salt Stress, and Secretion gene counts, and the second-highest Colonization count (**Figure 7**). Its elevated Salt Stress Tolerance gene count could be largely attributed to an abundance of heat shock protein genes, suggesting a probable physiological adaptation to salt stress (SM 10).

*B. thuringiensis* SRL368, with the most isolate-specific orthogroups also ranked high in Plant Hormone and VOC production genes (Fig. S8A; **Figure 7**), followed by SRL340 (**Figure 7**). The putative novel species SRL179 (second highest number of isolate-specific orthogroups) showed enrichment in Plant Colonization and Hormone/VOC genes, whereas the new species SRL337 (third highest number of isolate-specific orthogroups), had the fewest Osmoprotection and Secretion genes (Fig. S8A; **Figure 7**). *Paenibacillus* SRL342 and SRL398 carried many siderophore and iron-transport genes (Disease Resistance category), suggesting disease-protective potential for their host, and SRL342 also had the most Antioxidants and Spermidine synthesis genes (**Figure 7**; SM 10). These two isolates, both extracted from the same *Atriplex halimus* root sample, were isolated using different culture media (**Table 1**; SM 5). Their subtle differences may reflect microdiversity arising even within a single host microhabitat and/or culture-medium effects resulting from repeated subculturing of a diverse population of isolates of the same species in different media.

Interestingly, gene mining uncovered genes involved in the synthesis of enzymes and compounds associated with antifungal activity (e.g., cyclic lipopeptides such as surfactin, iturin, and fengycin, as well as chitinases), consistent with the antifungal effects observed in the *in vitro* dual-culture assays (**Figure 4**, SM 10). Gene mining further revealed a broad distribution among the 25 isolates of genes associated with VOCs biosynthesis: sulfur VOCs (methanethiol, dimethyl disulfide (DMDS), dimethyl trisulfide (DMTS)) known for their antifungal and biocontrol activities [96], and alcohol/ketone VOCs (acetoin, 2,3-butanediol, diacetyl) and their precursors, known for fungal pathogen inhibition and plant-growth promotion [97] (SM 10). These genomic features are consistent with the volatile-mediated antifungal effects observed in the dual-plate bioassays (**Figure 4**, Fig. S5).

### Concluding remarks

The genomic, phylogenetic, and functional characterization of 25 endophytic *Bacilli* isolates from plants with different lifestyles, revealed striking novelty across multiple levels. The phylogenomic analysis placed the isolates within a wide diversity of *Bacilli* lineages, spanning *Bacillus, Paenibacillus, Peribacillus, Neobacillus, Cytobacillus* and *Rossellomorea.* The reconstruction of the phylogenetic inference revealed a clear separation of several isolates, supporting the delineation of three novel species and expanding the known diversity of plant-associated *Bacilli*.

BGC mining uncovered 312 BGCs, nearly 60% of which were novel, highlighting an exceptional and unexplored reservoir of secondary metabolites. The high proportion of BGCs showing no or very low similarity to known clusters suggests that such unique and novel biosynthetic repertoires may constitute a distinctive adaptive feature of these endophytic *Bacilli* and could also indicate a distinctive trait of endophytes in general, facilitating their colonization and interaction with host plants. Similarly, genome mining identified the presence of diverse genes linked to colonization and stress tolerance reflecting ecological strategies that enable *Bacilli* to thrive in saline and nutrient-limited habitats. Bioassays validated the high-salinity tolerance, and strong antagonism against diverse phytopathogens, underscoring their potential as bioinoculants in agrifood.

Our study demonstrates that *Bacilli* endophytes from wild and halophytic plants combine taxonomic novelty with functional and biosynthetic innovation. Together, these findings underscore the dual importance of *Bacilli* endophytes: as promising resources for further investigation of their potential roles in crop resilience and biocontrol, and model systems for advancing our understanding of microbial adaptation, symbiosis, and the evolution of secondary metabolism.

## Acknowledgments

We thank Prof. A. M. Eren for his valuable help and insightful suggestions regarding the anvi’o pangenome analysis pipeline. *Ralstonia solanacearum* GMI1000 was kindly provided by Prof. Nemo Peeters (INRA CNRS Research Unit, France), *Clavibacter michiganensis* HMU4521 was kindly provided by Prof. Dimitris Goumas (Collection of the Department of Agriculture, Hellenic Mediterranean University, HMU), and *Paracidovorax citrulli* (formerly *Acidovorax citrulli*) ATCC29625 was kindly provided by Prof. Ronald Walcott (Department of Plant Pathology, University of Georgia).

## Availability of data and materials

Sequences and assemblies are registered at ENA with the PRJEB93985 accession number.

Genome and statistics analysis repository: https://github.com/nik-arapitsas/Bacillus_project

Phylogenetic analysis repository: https://github.com/FranziskaReden/bacillus_project

SM: Available at https://doi.org/10.5281/zenodo.18496603.

- Supplementary_Tables.docx
- Supplementary_Figures.pdf
- SM 1 = Supplementary_Material_1.docx
- SM 2 = Supplementary_Material_2_in_vitro_Antifungal_Data.xlsx
- SM 3 = Supplementary_Material_3_Sequencing_Data.xlsx
- SM 4 = Supplementary_Material_4_Checklist_GSC_MIxS.xlsx
- SM 5 = Supplementary_Material_5_Collective_Isolates_Data.xlsx
- SM 6 = Supplementary_Material_6_Identity_Matrix.xlsx
- SM 7 = Supplementary_Material_7_Gene_Lists_query_terms_with_refs.xlsx
- SM 8 = Supplementary_Material_8_OrthoFinder_statistics_per_isolate.xlsx
- SM 9 = Supplementary_Material_9_BGC_Data.xlsx
- SM 10 = Supplementary_Material_10_Genome_Mining_Data.xlsx

## Declarations

### Ethics approval and consent to participate

Not applicable.

## CRediT

Conceptualization: PFS, CAC

Data curation: SP, NPA, CAC, SS

Formal analysis: NPA, CAC, SP, SS, FR

Funding acquisition: PFS

Investigation: NPA, CAC, SP, SS, CP, EA

Methodology: CAC, NPA, SP, SS, FR

Project administration: PFS, SP, CAC

Resources: PFS, EAM, AS

Software: NPA, SP, FR, CAC

Supervision: PFS, EAM, AS

Validation: PFS, CAC, NPA, SP

Visualization: NPA, SP, FR, SS

Writing – original draft: NPA, PFS, SP, CAC, SS, FR, EAM, AS

Writing – review & editing: All authors reviewed and edited the manuscript

