## Supplementary Figures for "Unravelling the genomic and functional arsenal of *Bacilli* endophytes from plants with different lifestyles"

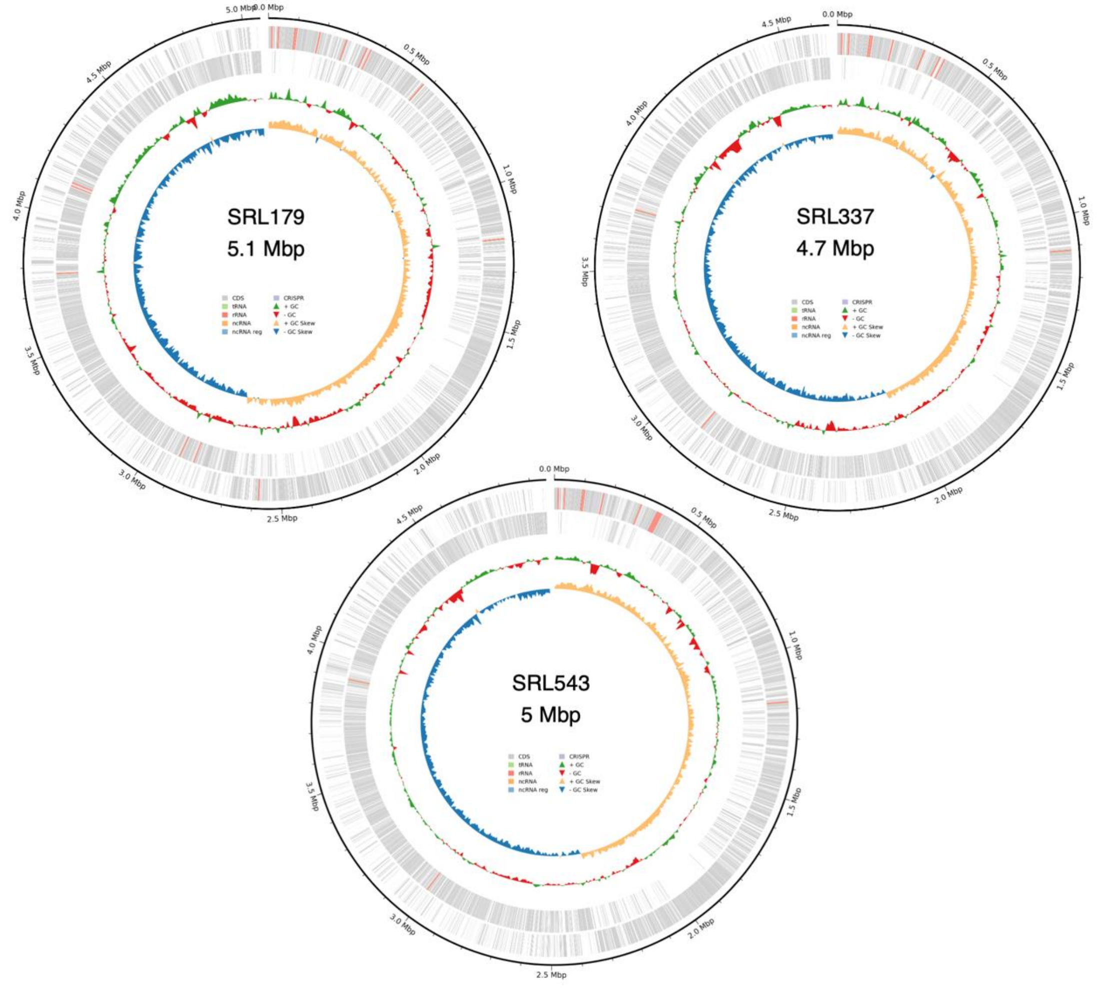


**Supplementary Figure S1** Whole-genome maps of the isolates SRL179, SRL337 and SRL543, representing putative new species, produced using the Bakta annotation pipeline.

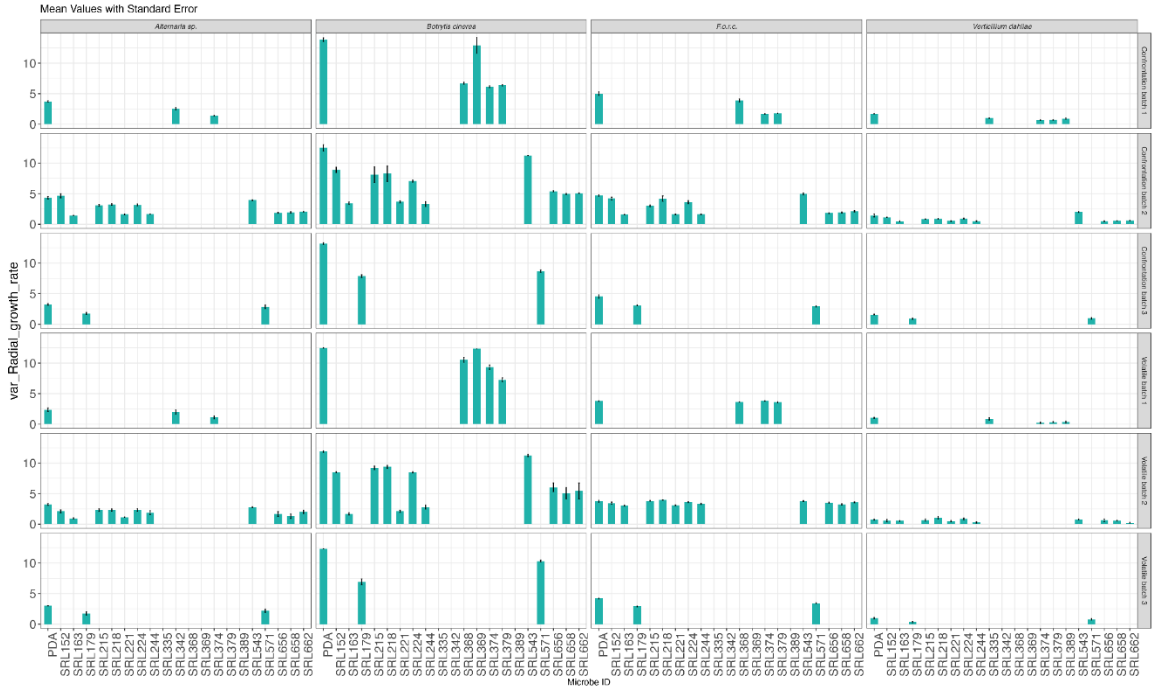


**Supplementary Figure S2** Bar plot for all *in vitro* tested fungal phytopathogens for the different batches and assays concerning the radial growth rate in mm.

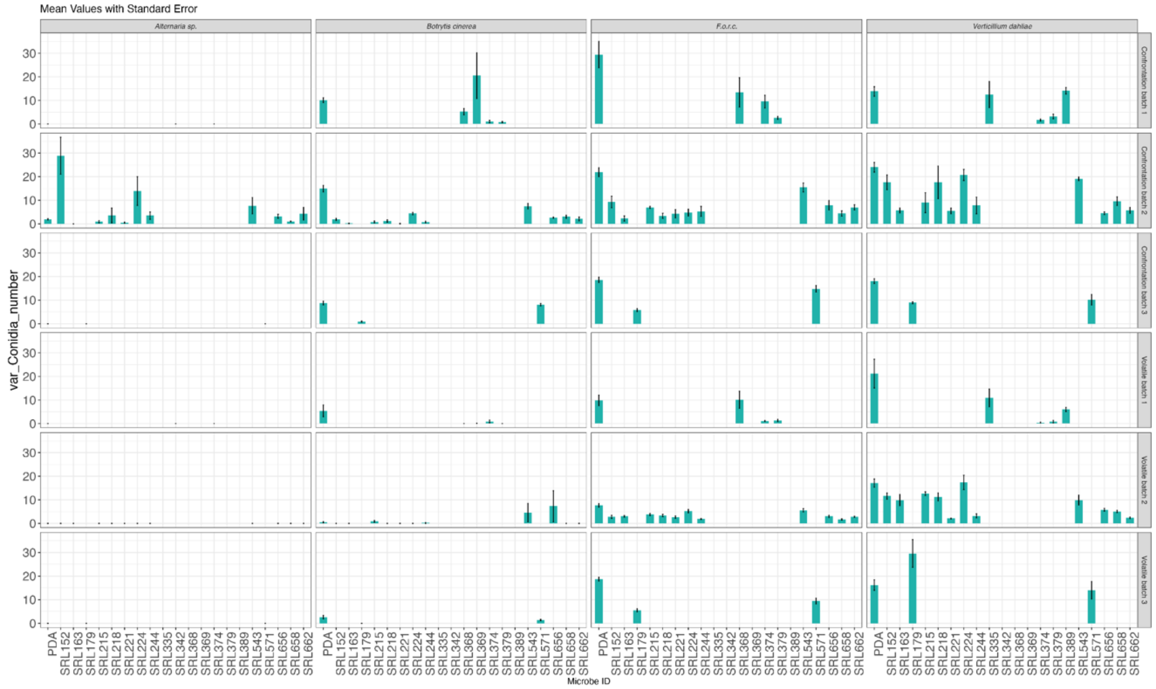


**Supplementary Figure S3** Bar plot for all *in vitro* tested fungal phytopathogens for the different batches and assays concerning the number of conidia.

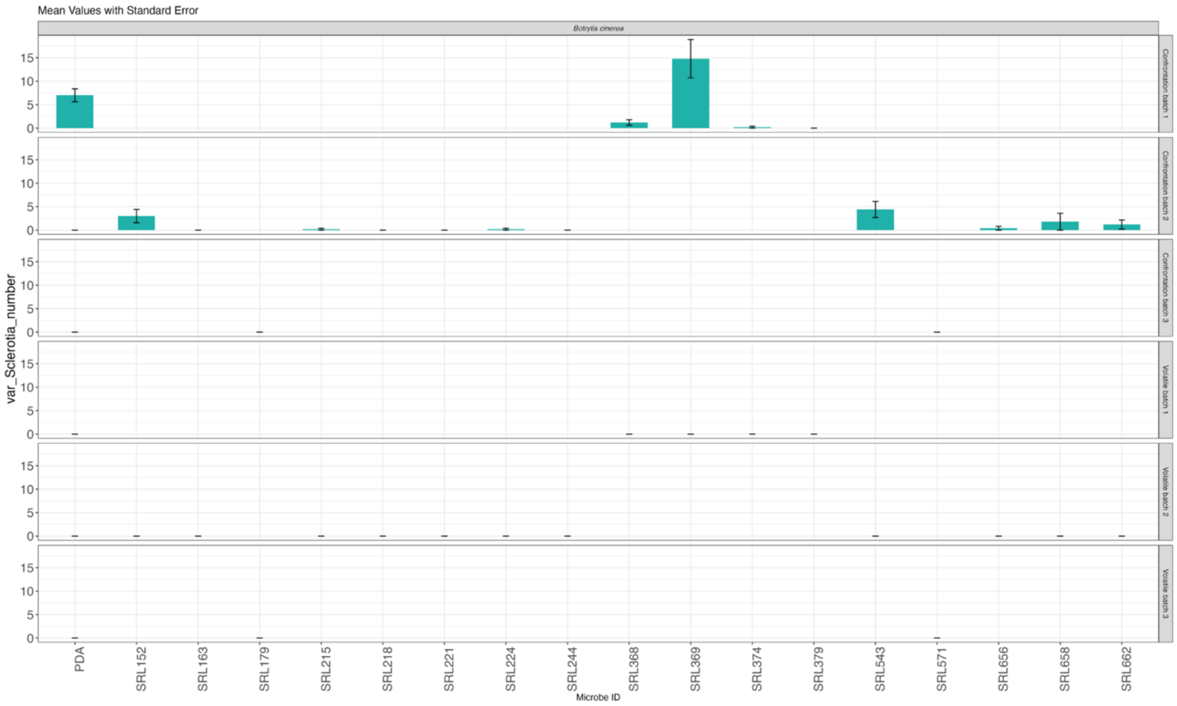


**Supplementary Figure S4** Bar plot for *Botrytis cinerea* pathogen for the different batches and assays concerning the number of sclerotia.

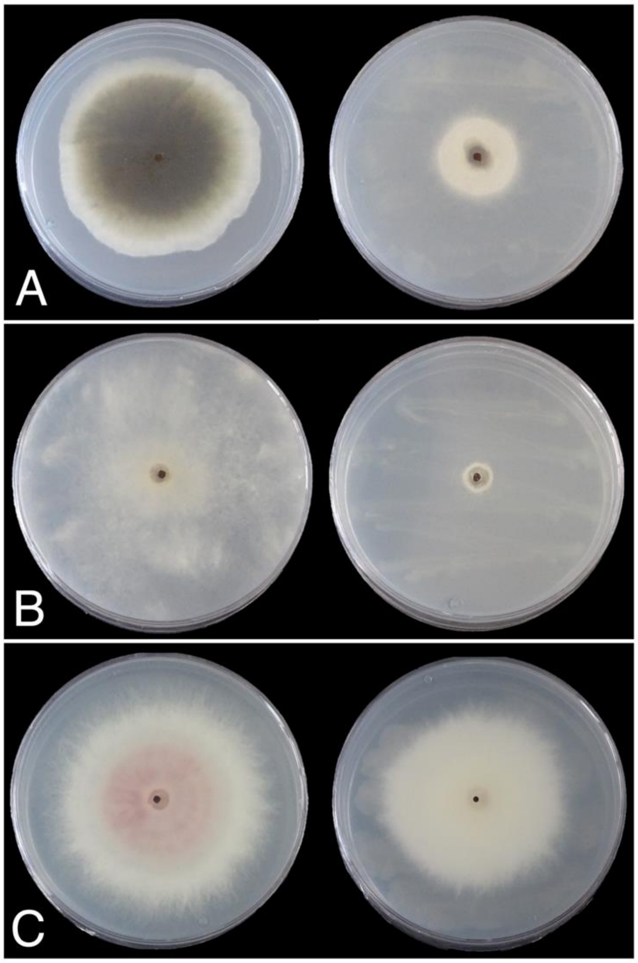


**Supplementary Figure S5** Inhibition of fungal growth of **A**. *Alternaria sp.*, **B**. *Botrytis cinerea* and **C**. *Fusarium oxysporum f.sp. radicis-cucumerinum* by the most effective *Bacillus* isolate (SRL163) in dual-plate assays.

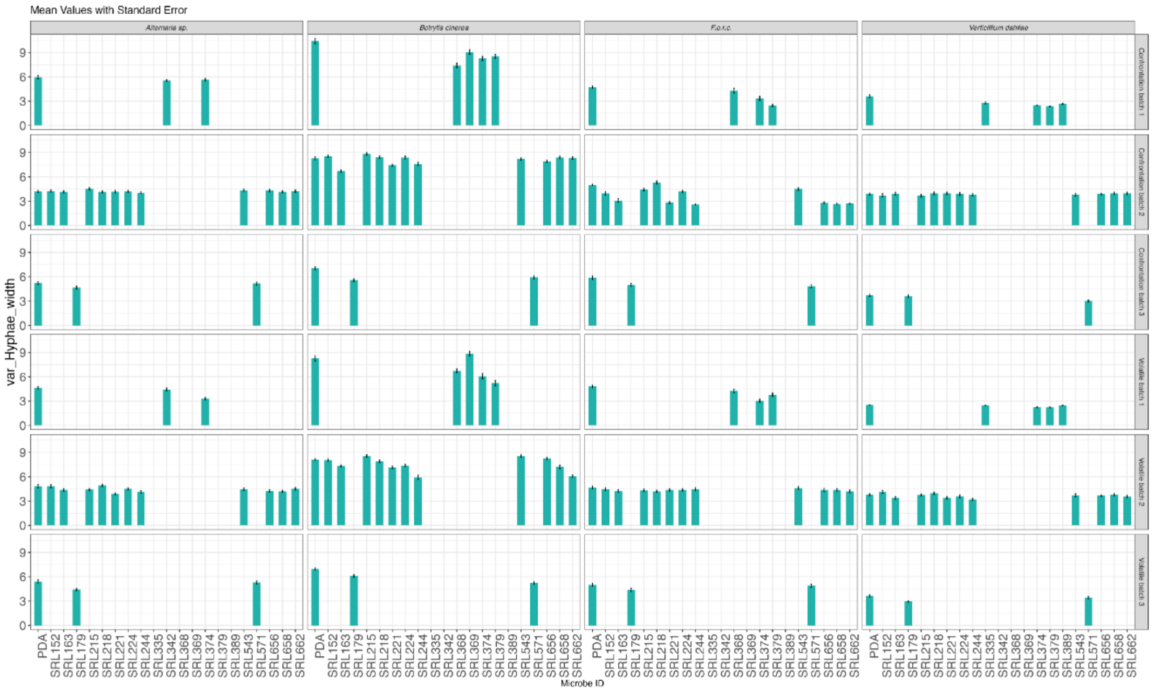


**Supplementary Figure S6** Bar plot for all *in vitro* tested fungal phytopathogens for the different batches and assays concerning the hyphae width measured in µm.

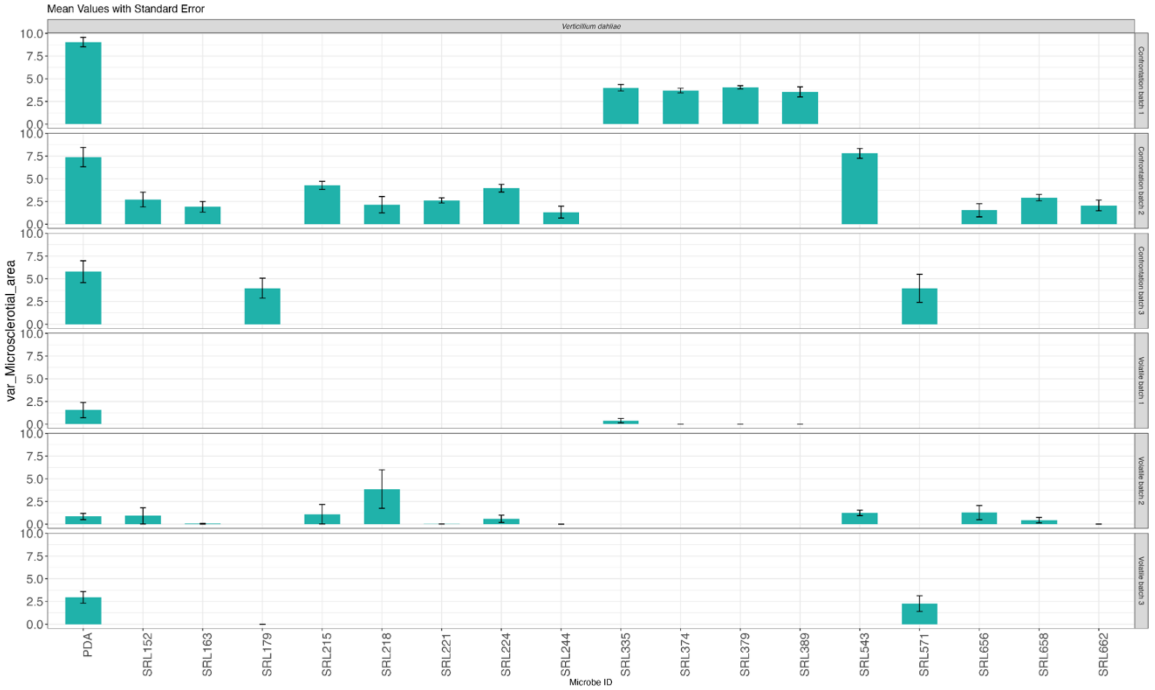


**Supplementary Figure S7** Bar plot for all *in vitro* batches for *Verticilium dahliae* concerning the microsclerotial area measured in $cm^2$.

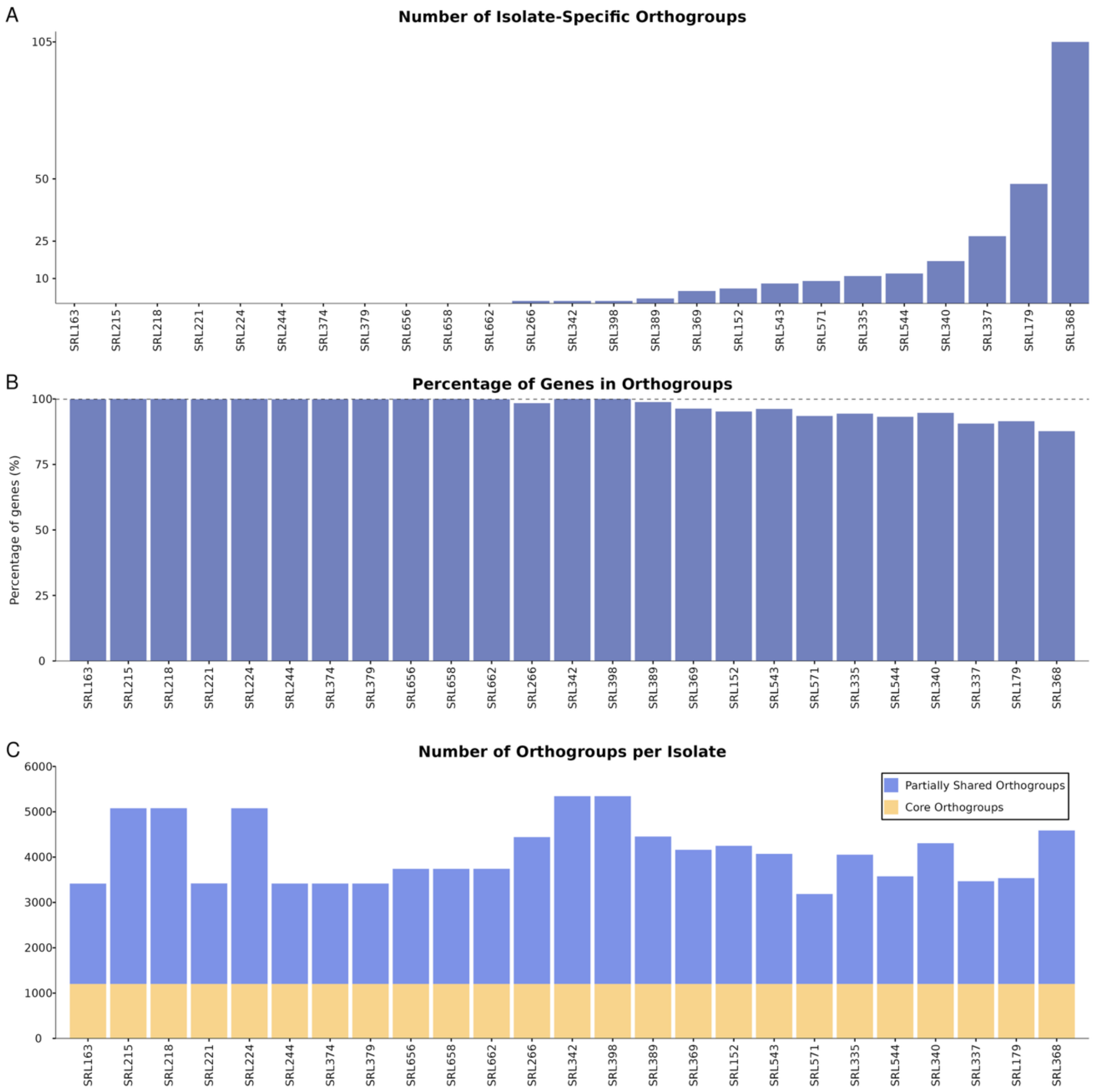


**Supplementary Figure S8 A.** Number of isolate-specific orthogroups per isolate. **B.** Percentage of genes attributed to orthogroups across isolates. **C.** Number of orthogroups per isolate shared with all isolates (core orthogroups) (yellow, bottom bars) and orthogroups shared with at least one but not all isolates (partially shared orthogroups) (blue, top bars).

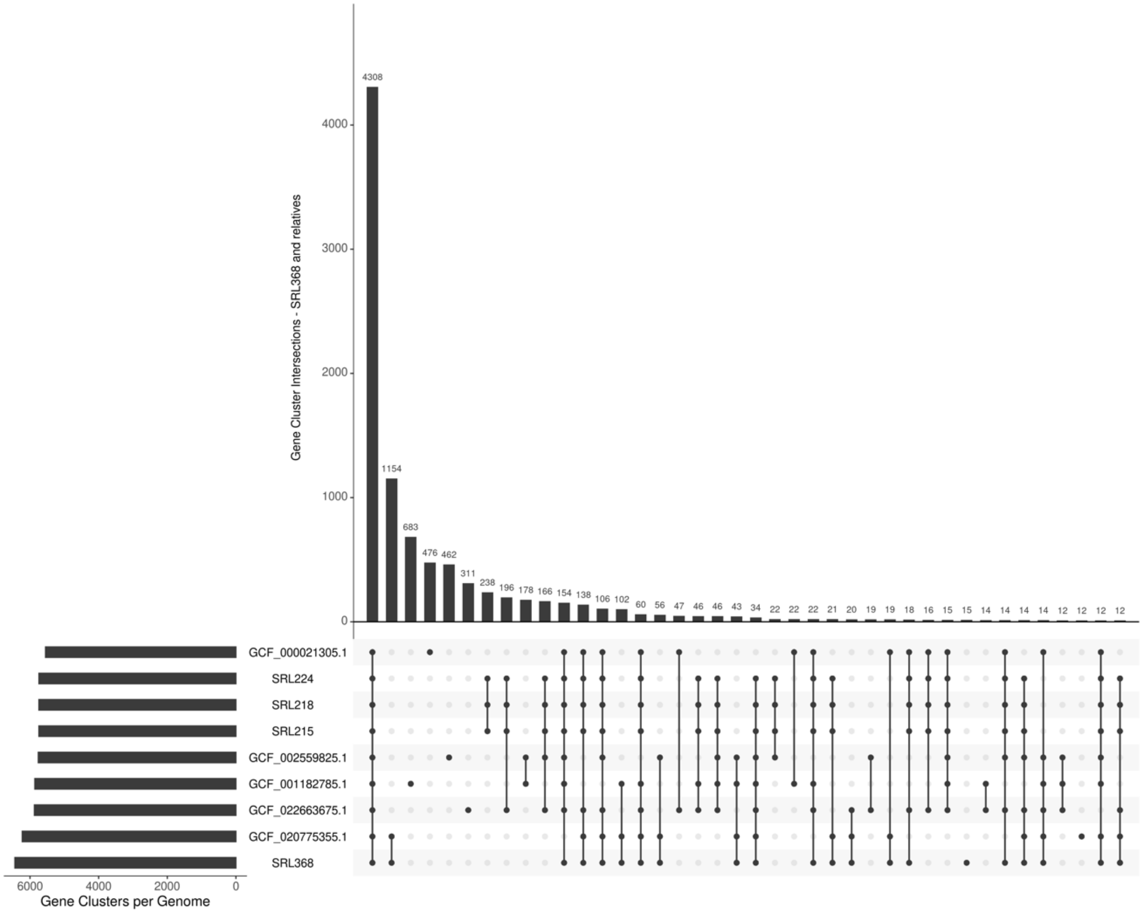


**Supplementary Figure S9** UpSet plot of the pangenome analysis of the *Bacillus thuringiensis* isolate SRL368 and its five closest relatives, including the *B. thuringiensis* isolates SRL215, SRL218 and SRL224. Horizontal bars (left) represent the total gene clusters per genome, and vertical bars (top) display the number of shared clusters for each intersection (genome combination) indicated by the connected dots below.

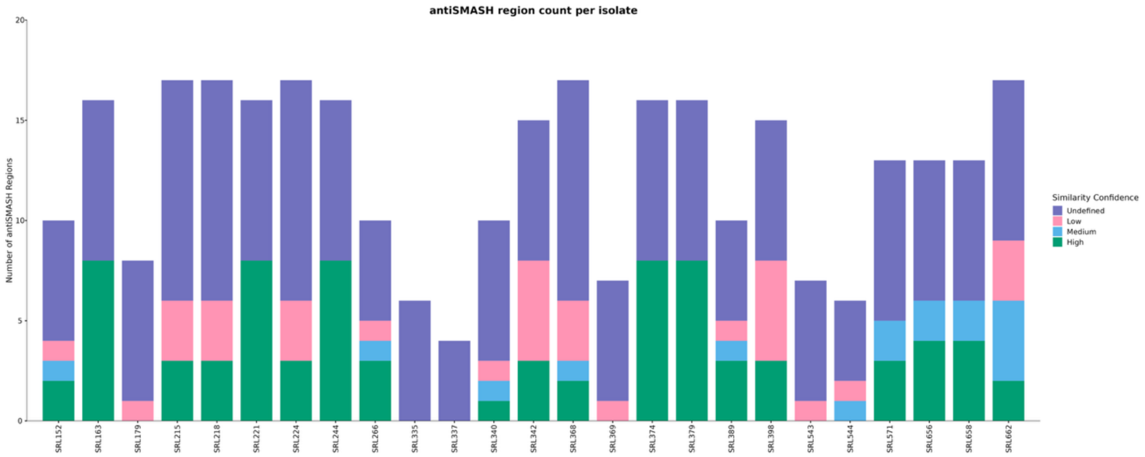


**Supplementary Figure S10 Distribution of BGC counts per isolate by similarity confidence**. Each stacked bar plot shows the number of BGCs identified in each isolate. Colors indicate the similarity confidence of each BGC to known clusters based on the antiSMASH database: High, Medium, Low, and Undefined (no close matches).

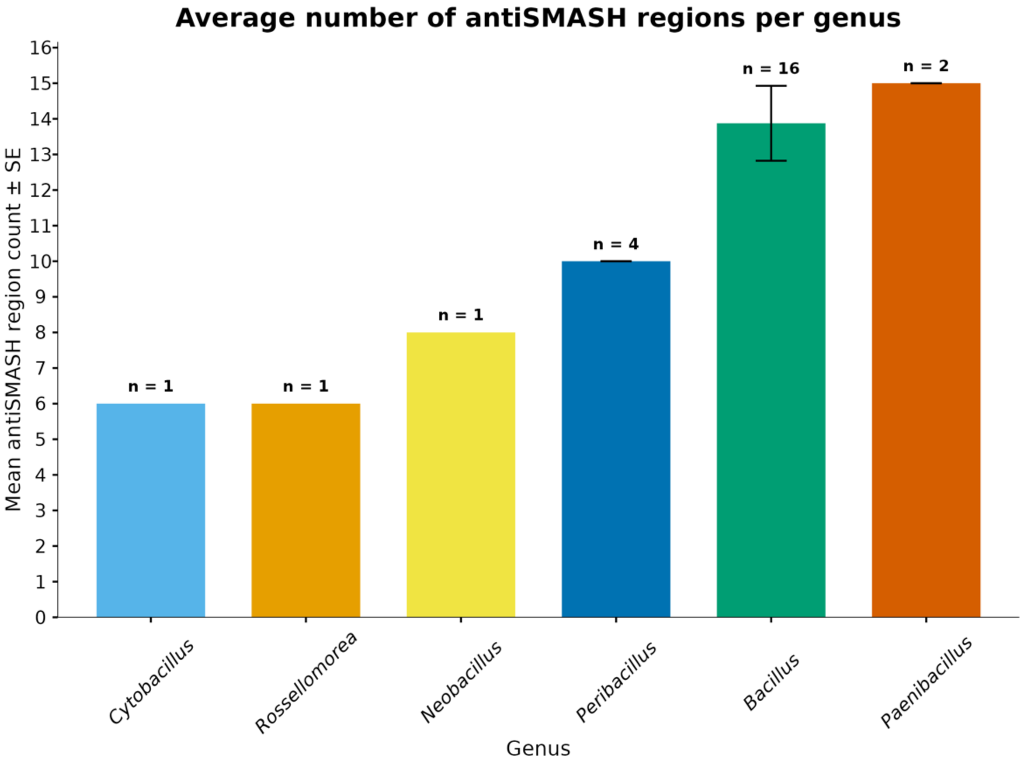


**Supplementary Figure S11 Average number of BGCs per genus across the genomes of the 25 isolates.** Bars showing mean antiSMASH region counts per genus ± standard error (SE). The value of n above each bar indicates the number of isolates, among the 25 analyzed, belonging to each genus.

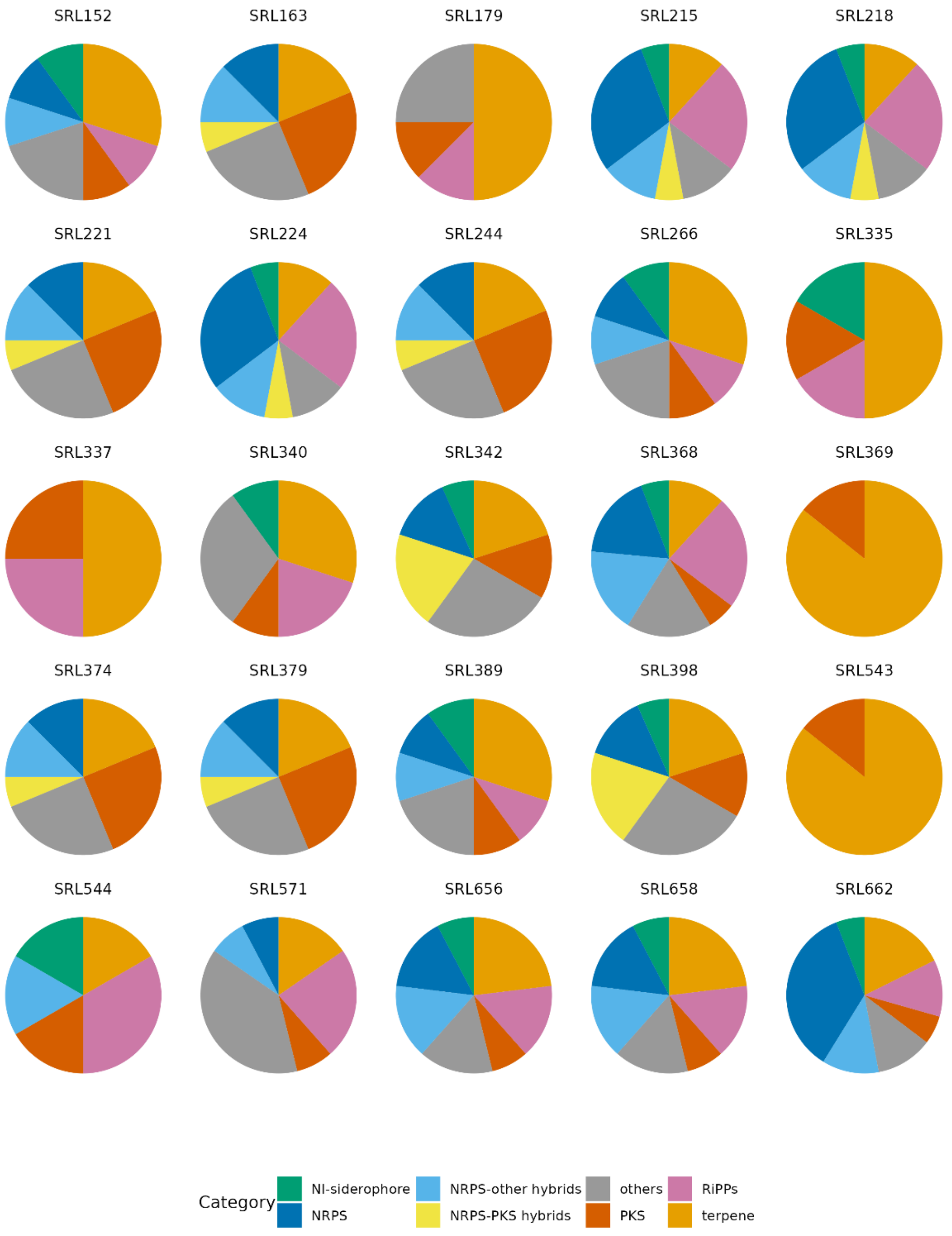


**Supplementary Figure S12 BGC classes content of each isolate.** Pie charts showing the relative proportions of BGC classes identified in each of the 25 isolates. Different colors correspond to eight BGC categories: NI-siderophores, NRPS, NRPS-PKS hybrids, NRPS-other hybrids, others, PKS, RiPPs and terpenes (see SM 1 - Supplementary Methods). For each isolate, percentages are calculated by dividing the number of BGCs of a given class by the total number of predicted BGCs in that isolate.
