## Supplementary Material 1 for "Unravelling the genomic and functional arsenal of *Bacilli* endophytes from plants with different lifestyles"

### Supplementary Methods

#### Detailed method for plant surface sterilization and endophytic bacterial isolation

Plant surface sterilization and endophytic bacterial isolation were performed as previously described [1]. Briefly, leaves and roots of selected samples were separated, washed gently with sterile distilled water to remove soil and dust particles, dried using clean absorbent paper and weighted on a precision laboratory balance. For surface sterilization, plant materials were placed in ethanol 75% v/v for 60 s with shaking, then in sodium hypochlorite solution 3% w/v (NaClO) for 10 min, and finally in ethanol 75% v/v for 60 s. The NaClO was removed completely by rinsing plant materials at least 10 times with sterile distilled water (dH_2_O). Part of the last rinse (100 μL) was plated on three different solid media: Nutrient Agar (NA) (5 g/L bacto-peptone, 3 g/L yeast extract, 5 g/L NaCl, 15 g/L agar, pH 7.4), Reasoner's 2A (R2A) agar (0.5 g/L yeast extract, 0.5 g/L proteose peptone, 0.5 g/L casamino acids, 0.5 g/L glucose, 0.5 g/L soluble starch, 0.3 g/L sodium-pyruvate, 0.3 g/L K_2_PO_4_, 0.05 g/L MgSO_4_·7H_2_O, 15 g/L agar, pH 7.2) and NA1/2 (2.5 g/L bacto-peptone, 1.5 g/L yeast extract, 2.5 g/L NaCl, 15 g/L agar, pH 7.4), and monitored for microbial growth to evaluate surface sterilization efficiency. Only successfully sterilized plant material was processed further. The surface sterilized roots and leaves from each sample were cut thoroughly using sterile tongs and scissors. Sterile 10 mM MgSO_4_ solution was added in the slashed tissue in a ratio of 10 mL solution per 1 g of roots or leaves, and the tissue was ground into a slurry using an autoclaved mortar and pestle. The slurry was transferred in sterile 50 mL tubes and placed onto a rotary shaker, spinning at 150 rpm, at 25 °C for 2 h. Following the shaking process, 50 µL of the surface-sterilized root or leaf extract was plated in triplicate onto NA, R2A and NA1/2 plates and incubated at 28°C for 4 - 7 days. Occasionally, the plates were checked for growth. Distinct single colonies were selected based on their colour, texture and morphology and repeatedly streaked to obtain pure cultures. Colonies of each pure isolate were dissolved in their respective liquid medium in 50% v/v glycerol and stored at −80 °C. Isolate SRL340 was obtained from R2A and SRL342 from NA1/2 medium. The rest of the isolates were obtained from NA plates. The culture of the isolates was continued in the same media from which they were firstly isolated.

#### Identification of isolates with 16S rRNA gene sequencing

For preliminary identification, colonies were suspended in 100 μL sterile dH_2_O, heated at 98 °C for 10 min and centrifuged at 10,000 × g for 10 min. For 16S rRNA gene amplification, 2 μL of the supernatant were mixed with 1× Taq Pol Buffer (Minotech, Heraklion, Greece), 0.5 μM of primer 27F (5′-AGAGTTTGATCCTGGCTCAG-3′) [2], 0.5 μM of primer 1492R (5′-GGTTACCTTGTTACGACTT-3') [3], 0.2 mM dNTPs and 0.04 U Taq DNA Pol (Minotech, Heraklion, Greece) in 25 μL reactions. PCR was performed on a BioRad T-100 Thermocycler at 94◦C for 2 min, followed by 30 cycles of 45 s at 94◦C, 30 s at 55◦C, 90 s at 72◦C, and a final extension at 72◦C for 10 min. Products were puriﬁed using NucleoSpin® Gel and PCR Clean-up kit (Macherey-Nagel™, Düren, Germany) according to the manufacturer’s instructions and sequenced by Macrogen (Europe) using primer 27F. The resulting chromatograms were quality-checked and manually trimmed in BioEdit [4], and resulting sequences were queried and identified with BLASTN [5]. The identification results of the 25 isolates based on the 16S rRNA gene sequencing are provided in the SM 5.

#### Detailed method for bacterial whole genome extraction for next-generation sequencing

The isolation of total genomic DNA was based on a protocol by A. Kiledal and J. A. Maresca [6]. Each bacterial isolate was grown in its respective medium (NA, R2A or NA1/2) and a single bacterial colony was used to prepare a 10 mL liquid culture with NB (Nutrient Broth) (NA medium without agar), NB1/2 or R2A medium without agar accordingly. After incubation at 28°C, 200 rpm for 16 h, the cultures obtained an OD_600_ of approximately 0.8 - 1.2 and centrifuged at 5000 rpm for 10 min at 28°C. Cells were re-solubilized in 400 μL of TEN solution (10 mM Tris-HCl pH 8.0, 10 mM EDTA, 150 mM NaCl). Cell wall was disrupted with 20 μL of lysozyme (20 mg/mL in ultrapure water) (Sigma Aldrich, St. Louis, Missouri, United States) followed by incubation at 37°C for 20 min. RNA was removed by the addition of 2 μL of RNase A (20 mg/mL) (Macherey-Nagel™, Düren, Germany) and incubation at 65°C for 3 min. For disruption of the cell membrane and protein degradation, 40 μL SDS (10% w/v), 10 μL of proteinase K (10 mg/mL) (Sigma Aldrich, St. Louis, Missouri, United States) and 550 μL of TEN* solution (10 mM Tris-HCl pH 8.0, 1 mM EDTA, 50 mM NaCl) were added, followed by incubation at 60°C for 2 h with occasional agitation. To avoid the interference of possible surfactants produced by the isolates with the phase separation, 0.1 volume of 3 M sodium acetate and 1 volume of cold isopropanol were used and the samples were incubated at 4°C for 20 min. Then, followed centrifugation at 5000 rpm for 10 min, discard of the supernatant and resuspension of the pellet in a mix of 400 μL TEN and 550 μL TEN*. The suspension was mixed with 900 μL of phenol. Followed by centrifugation at 16,000 × g for 5 min, the aqueous phase was separated. The extraction was repeated once with 900 μL of phenol and twice with 900 μL of chloroform and isoamyl alcohol solution in a 24:1 ratio. The aqueous phase of the last separation was transferred in 10 mL -20°C absolute ethanol. After a mild mix, the DNA formed profound threads in the ethanol that were coiled up with the bended tip of a sterile glass Pasteur pipette and air dried under a fume hood for 10 min. Then, the tip of the Pasteur pipette was inserted in a 1.5 mL microtube and the dried DNA was dissolved in 100 μL 10 mM Tris-HCl pH 8.0 solution and stored at -20°C. DNA size and quality were estimated with agarose gel electrophoresis and DNA was quantified with NanoDrop™ Spectrophotometer (Thermo Fisher Scientific Inc., Waltham, Massachusetts, United States) and Qubit fluorometer (Thermo Fisher Scientific Inc., Waltham, Massachusetts, United States). DNA samples were sent to commercial vendor Novogene (Netherlands) for Illumina NovaSeq 2 x150 bp sequencing, and PacBio Sequel II long read sequencing.

**Detailed method for phylogenetic tree construction**

Taxonomic assignments were conducted with GTDB-tk [7], and FastANI [8]. For genomes with Average Nucleotide Identity (ANI) ≤ 95%, we used the *de novo* GTDB-tk workflow to infer trees that contained our genomes and the GTDB-Tk reference genomes of the Bacillota phylum. Contigs below 200 bp were removed from the assemblies.

A species tree was then inferred from the same 120 marker genes used in the GTDB-Tk pipeline, including all 25 assemblies analysed in this study. Unlike GTDB-Tk, which relies on a subset of sites, full-length marker genes were employed. First, gene annotation was performed with Prokka v1.14.6 [9] using default parameters to ensure consistent and reproducible annotation across all genome assemblies. This includes using Prokka’s default protein databases, which prioritize UniProt curated entries, as well as the default BLASTp e-value cutoff of 1e-6 with a minimum query coverage of 80%. Then, the 120 unique marker genes used in the GTDB-Tk pipeline were extracted from assemblies.

Next, the 120 full-length marker genes used to construct the GTDB bacterial reference tree were downloaded from the GTDB website [10]. From this set, the marker genes of reference genomes belonging to the same putative genera as the query assemblies were selected based on the GTDB taxonomy. Marker genes from non-reference GTDB genomes were included when they represented the closest putative relatives. These closest putative relatives were determined by calculating the average relative sequence identity across pairwise aligned marker genes between GTDB genomes and query assemblies within the same genus. The genome with the smallest distance was retained, irrespective of its reference status. To avoid redundancy, highly similar reference genomes were removed from further analysis. Reference genomes showing an average relative marker gene identity greater than 95% (see identity matrix, SM 6) were grouped, and only one genome per group was retained for downstream analysis, resulting in a final set of 280 GTDB reference genomes. Protein sequences of marker genes were aligned from the query assemblies and the set of 280 GTDB reference genomes via MUSCLE v5.1 [11]. Initial alignments were inferred using the stratified option to generate an ensemble of 16 entropy-stratified alignments to consider alignment uncertainty. The resulting amino acid alignment with the highest overall column confidence was then used as a template to project gaps back onto the corresponding coding DNA sequences, thereby generating codon-aware DNA alignments that preserve the correct reading frame.

Gene trees were inferred on each codon-aware DNA alignment using IQ-TREE 2 v2.4.0 [12]. The best-fit nucleotide substitution model for each gene was determined using ModelFinder [13] as implemented in IQ-TREE. To account for phylogenetic uncertainty and provide a basis for downstream consensus and support analysis, 50 independent maximum-likelihood (ML) trees were inferred per alignment. Tree inferences were executed with the –keep-ident flag enabled to retain potentially identical genes of highly similar genomes during the analysis. Topology support was evaluated using the approximately unbiased (AU) [14] test and other tests implemented in IQ-TREE [12]. All trees were retained for further analysis, as each showed statistically significant support in at least one test.

Consequently, the full set of 6,000 ML gene trees (120 marker genes × 50 trees) was used as input for species tree inference in ASTRAL-IV [15]. For the ASTRAL analysis, 16 subsampling rounds were specified for both the exploration and initial subsampling steps. Furthermore, the average gene length parameter was set to 1300 bp, which corresponds to the mean length of the marker genes in the studied dataset and allows ASTRAL to scale branch lengths appropriately. Thereafter, the ballistic distances—defined as the sum of branch lengths along the unique path between two tips—were calculated in the species tree between each assembly and its closest putative relative. Species delimitation was subsequently performed on the species tree using mPTP v0.2.5 [16] with maximum likelihood optimization in single-rate mode. Phylogenetic trees were visualized with TreeViewer [17]. All analyses were executed through a Snakemake workflow available on GitHub (<https://github.com/FranziskaReden/bacillus_project_phylogeny.git>) under the Creative Commons license (CC0 1.0).

**BGCs categories**

The BGCs were organized in the following categories:

• **Non-ribosomal peptide synthetases (NRPS)**: NRPS and NRPS-like BGCs

• **Terpenes**: terpenes and terpene-precursors

• **Polyketide synthases (PKS)**: T3PKS, transAT-PKS, PKS-like BGCs and HR-T2PKS

• **NRPS-independent IucA/IucC-like siderophores (****NI-siderophores)**: NI-siderophores

• **NRPS-PKS hybrids**: BGCs that combine elements of both NRPS and PKS e.g. NRPS.T1PKS.transAT-PKS

• **Ribosomally synthesized and post-translationally modified peptide products (RiPPs)**: RiPP-like BGCs and azole-containing-RiPPs

• **NRPS-other hybrids**: BGCs that combine elements of NRPS and other non-PKS BGCs, e.g. NRPS.betalactone

• **Others**: the rest BGCs which does not fit in the categories above e.g. lassopeptides, lanthipeptides, phosphonates, etc.
