## Supplementary Tables for "Unravelling the genomic and functional arsenal of *Bacilli* endophytes from plants with different lifestyles"

**Supplementary Table S1**. Geographic coordinates and host plant species of samples collected across Crete and Chrysi island.

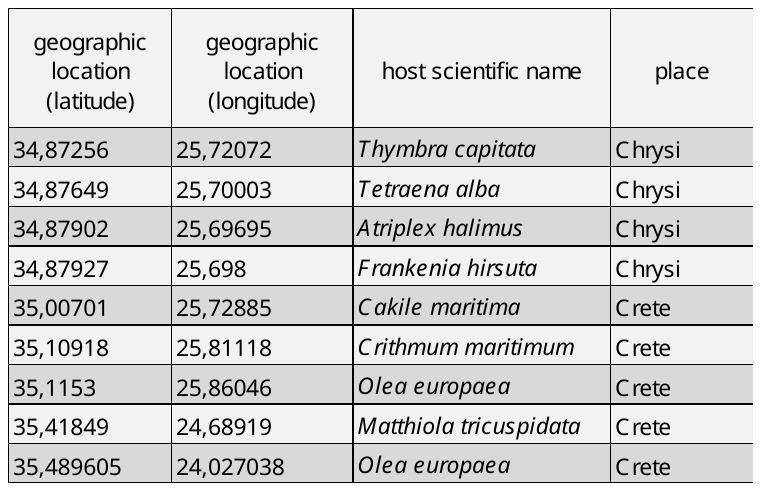

**Supplementary Table S2.** Summary of the *in vitro* experiments against fungal phytopathogens.

| **pathogen** | **type_bioassay** | **microbes** | **microbe_id** | **Petri #** |
| --- | --- | --- | --- | --- |
| *Alternaria* sp. | Confrontation | 15 | SRL342,SRL374,SRL662,SRL656,SRL244,SRL152,SRL215,SRL218,SRL224,SRL543,SRL221,SRL163,SRL658,SRL179,SRL571 | 90 |
| *Alternaria* sp. | Volatile | 15 | SRL342,SRL374,SRL662,SRL656,SRL244,SRL152,SRL215,SRL218,SRL224,SRL543,SRL221,SRL163,SRL658,SRL179,SRL571 | 90 |
| *Botrytis cinerea* | Confrontation | 17 | SRL368,SRL369,SRL374,SRL379,SRL662,SRL656,SRL244,SRL152,SRL215,SRL218,SRL224,SRL543,SRL221,SRL163,SRL658,SRL179,SRL571 | 100 |
| *Botrytis cinerea* | Volatile | 17 | SRL368,SRL369,SRL374,SRL379,SRL662,SRL656,SRL244,SRL152,SRL215,SRL218,SRL224,SRL543,SRL221,SRL163,SRL658,SRL179,SRL571 | 100 |
| *F.o.r.c.* | Confrontation | 16 | SRL368,SRL374,SRL379,SRL662,SRL656,SRL244,SRL152,SRL215,SRL218,SRL224,SRL543,SRL221,SRL163,SRL658,SRL179,SRL571 | 95 |
| *F.o.r.c.* | Volatile | 16 | SRL368,SRL374,SRL379,SRL662,SRL656,SRL244,SRL152,SRL215,SRL218,SRL224,SRL543,SRL221,SRL163,SRL658,SRL179,SRL571 | 95 |
| *Verticillium dahliae* | Confrontation | 17 | SRL335,SRL374,SRL379,SRL389,SRL662,SRL656,SRL244,SRL152,SRL215,SRL218,SRL224,SRL543,SRL221,SRL163,SRL658,SRL179,SRL571 | 100 |
| *Verticillium dahliae* | Volatile | 17 | SRL335,SRL374,SRL379,SRL389,SRL662,SRL656,SRL244,SRL152,SRL215,SRL218,SRL224,SRL543,SRL221,SRL163,SRL658,SRL179,SRL571 | 100 |

**Supplementary Table S3.** Overall OrthoFinder statistics for the genomes of the 25 isolates.

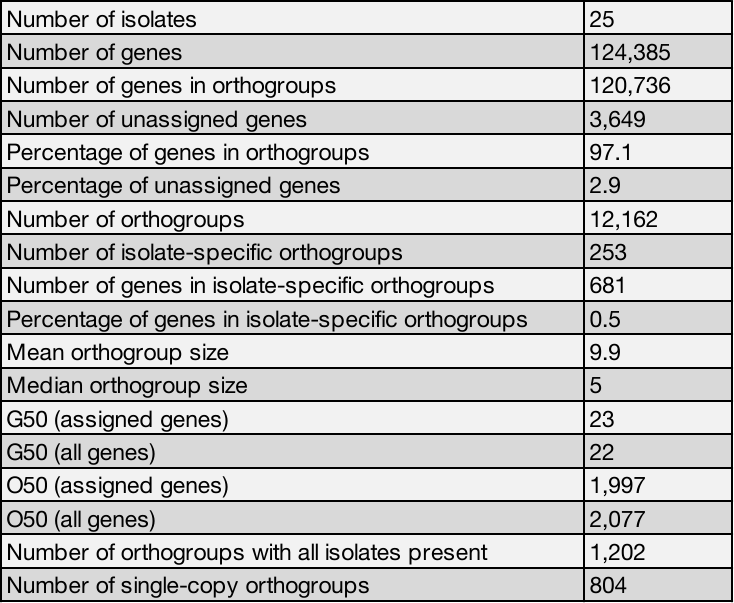
